# Chromatin interaction aware gene regulatory modeling with graph attention networks

**DOI:** 10.1101/2021.03.31.437978

**Authors:** Alireza Karbalayghareh, Merve Sahin, Christina S. Leslie

## Abstract

Linking distal enhancers to genes and modeling their impact on target gene expression are longstanding unresolved problems in regulatory genomics and critical for interpreting non-coding genetic variation. Here we present a new deep learning approach called GraphReg that exploits 3D interactions from chromosome conformation capture assays in order to predict gene expression from 1D epigenomic data or genomic DNA sequence. By using graph attention networks to exploit the connectivity of distal elements up to 2Mb away in the genome, GraphReg more faithfully models gene regulation and more accurately predicts gene expression levels than state-of-the-art deep learning methods for this task. Feature attribution used with GraphReg accurately identifies functional enhancers of genes, as validated by CRISPRi-FlowFISH and TAP-seq assays, outperforming both CNNs and the recently proposed Activity-by-Contact model. Sequence-based GraphReg also accurately predicts direct transcription factor (TF) targets as validated by CRISPRi TF knockout experiments via in silico ablation of TF binding motifs. GraphReg therefore represents an important advance in modeling the regulatory impact of epigenomic and sequence elements.

## Introduction

Transcriptional gene regulation involves the binding of transcription factors (TFs) at both promoter and enhancer regions and the physical interaction of these bound complexes via DNA looping. Technological advances in chromosome conformation capture assays such as Hi-C (Lieberman-Aiden et al., 2009), HiChIP (Mumbach et al., 2016), and Micro-C (Krietenstein et al., 2020) provide high-resolution data on 3D chromatin interactions, including regulatory interactions between enhancers and promoters. 3D interaction data sets combined with traditional 1D epigenomic profiles, including chromatin accessibility (DNase-seq and ATAC-seq (Buenrostro et al., 2015)) and histone modifications (via ChIP-seq and CUT&RUN (Skene and Henikoff, 2017)), together map the chromatin state and connectivity of regulatory elements and should provide rich training data for predictive gene regulatory models, including models that also incorporate the underlying genomic DNA sequence. Ultimately, such models could be used to infer the regulatory function of non-coding genetic variants in a cell type of interest.

Here we propose a gene regulatory modeling approach called GraphReg that integrates 1D epigenomic data (Epi-GraphReg) or both epigenomic and DNA sequence data (Seq-GraphReg) with 3D chromatin interaction data from HiChIP, Hi-C, or Micro-C via a graph neural network to predict gene expression. The 1D input data can include any standard epigenomic assays such as histone modification ChIP-seq, transcription factor ChIP-seq, or chromatin accessibility from DNase-seq or ATAC-seq.

GraphReg models use convolutional neural network (CNN) layers to learn local representations from 1D inputs, followed by graph attention network (GAT) layers to propagate these representations over the 3D interaction graph, in order to predict gene expression (CAGE-seq) across genomic positions (bins). GraphReg is trained to predict CAGE-seq (Shiraki et al., 2003), a tag-based protocol for gene expression measurement and transcription start site (TSS) mapping, since it quantifies promoter output and does not depend on transcript length.

Our motivation for proposing GraphReg is two-fold. First we aim to improve the accuracy of predictive gene regulatory models by leveraging 3D genomic architecture to incorporate distal enhancer elements, and we demonstrate that GraphReg outperforms baseline CNN models for prediction of CAGE output. Second, we seek to interpret the model to assess the functional importance of distal enhancers for regulation of specific target genes. To this end, we use feature attribution methods on trained GraphReg models and show that we obtain a better ranking of functional enhancers as validated by CRISPRi-FlowFISH (Fulco et al., 2019) and TAP-seq (Schraivogel et al., 2020) than baseline CNN models and the recently proposed Activity-by-Contact (ABC) model (Fulco et al., 2019). Finally, we demonstrate that Seq-GraphReg models capture meaningful TF binding signals through in silico motif ablation experiments, which accurately predict direct TF targets as validated by CRISPRi-based TF perturbation experiments.

## Results

### GraphReg: a deep graph neural network model for interaction aware gene expression prediction

In our experiments, we used a minimal set of 1D epigenomic data relevant to gene regulation: DNase-seq as a measure of chromatin accessibility, H3K4me3 ChIP-seq for promoter activity, and H3K27ac ChIP-seq for enhancer activity. For 3D interaction data, we have used a variety of chromatin conformation assays, such as Hi-C, H3K27ac HiChIP, and Micro-C. Since Epi-GraphReg is trained on 1D epigenomic and 3D interaction input data in a given cell type, in order to predict gene expression output in the same cell type, it learns a regulatory model that can generalize to other contexts. That is, given cell-type specific input data in a new cell type, the trained Epi-GraphReg model can predict gene expression in that cell type. In this sense, the Epi-GraphReg model is *cell-type agnostic*. Seq-GraphReg uses DNA sequence as the input and performs multi-task learning, predicting DNase, H3K4me3, and H3K27ac in the CNN block and predicting CAGE-seq after the GAT block. Seq-GraphReg is therefore a *cell-type restricted* model because it learns the TF binding signals specific to the training cell type and consequently cannot generalize to another cell type. However, Seq-GraphReg can capture cell-type specific TF binding motifs in enhancer and promoter regions and potentially predict the impact of DNA sequence alterations on target gene expression.

We bin the epigenomic data (H3K4me3, K3K27ac, and DNase) at 100bp resolution. We process the 5Kb resolution Hi-C/HiChIP/Micro-C contact matrices with the HiC-DC+ package (Carty et al., 2017; Sahin et al., 2021) to identify significant interactions of genomic distance up to 2Mb, and we use the 3D interactions satisfying three different false discovery rates (FDR) of 0.1, 0.01, and 0.001 to define a graph on 5Kb genomic bins. We process 3D interaction data up to 2Mb for two main reasons. First, the read coverage of 3D assays become sparser after 2Mb, making it difficult to robustly find significant interactions. Second, the majority of known functional enhancers of genes reside within 2Mb of the TSS. For consistency with the 3D interaction data, we also bin CAGE-seq at 5Kb resolution. In each batch of training, we extract 6Mb genomic regions and use the corresponding DNA sequences, epigenomic data, interaction graphs from 3D data, and the corresponding CAGE-seq as training examples for the model. For the next batch, we shift the entire 6Mb region by 2Mb and repeat this to cover the training chromosomes.

For Epi-GraphReg (**Fig. 1A**, **Supplemental Fig. S1**), each epigenomic signal track is treated as a different channel and fed to 1D CNN layers followed by max-pooling and ReLU activation functions. This produces latent local representations at 5Kb resolution, which serve as the node features for the interaction graphs derived from 3D data. We then use several graph attention network (GAT) layers to propagate these local representations over the interaction graphs, so that promoter regions are influenced by the representations of their candidate distal enhancers. Finally, we predict the CAGE values (promoter activities) by a fully connected (FC) layer.

**Figure 1:**
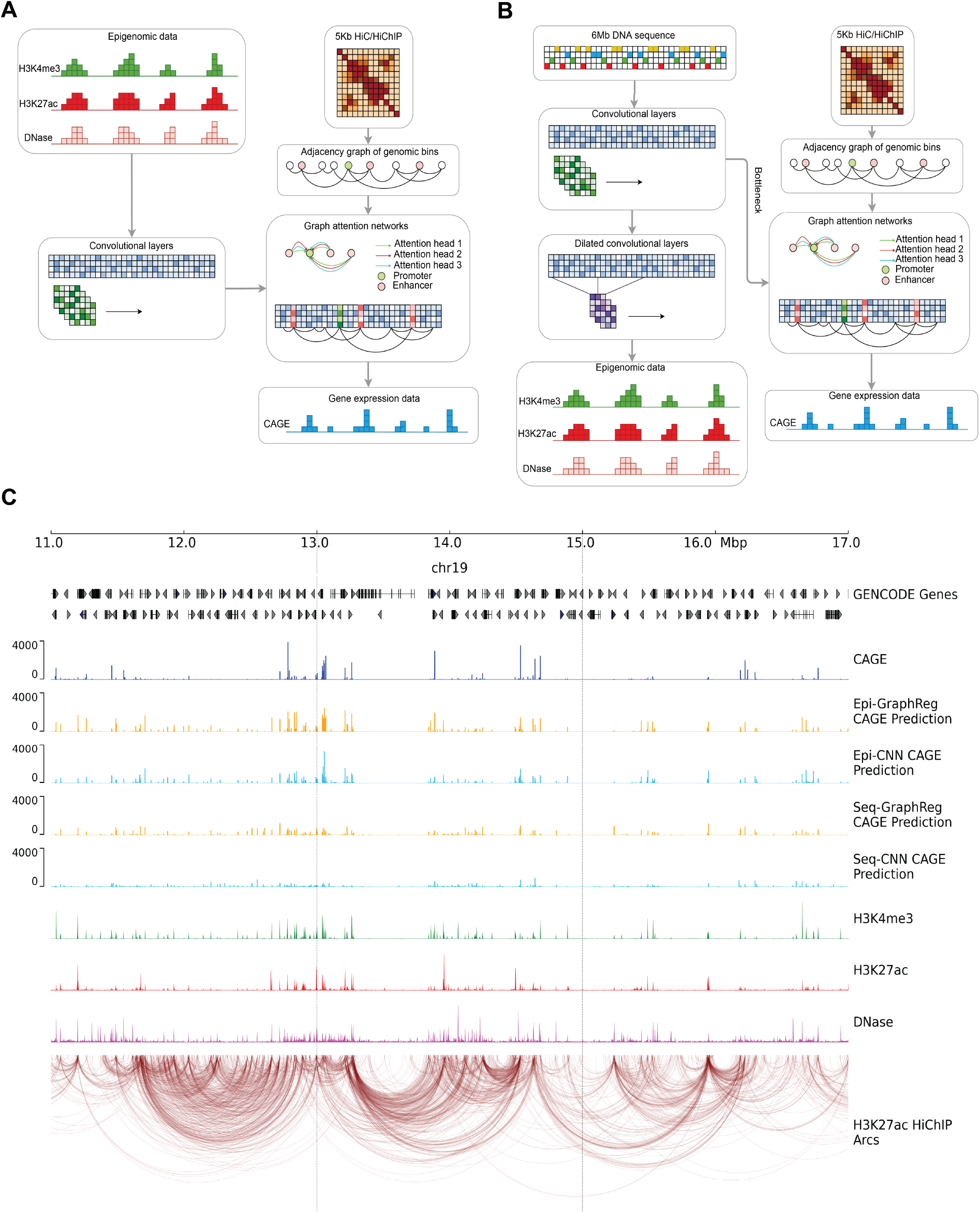
A schematic overview of GraphReg models. **A**. The Epi-GraphReg model uses 1D epigenomic data, such as H3K4me3 and H3K27ac ChIP-seq and DNase-seq (or ATAC-seq) to learn local features of genomic bins via convolutional neural networks, and then propagates these features over adjacency graphs extracted from HiC/HiChIP contact matrices using graph attention networks, in order to predict gene expression (CAGE-seq) across genomic bins. **B**. The Seq-GraphReg model uses DNA sequence as input, and after some convolutional and dilated convolutional layers predicts epigenomic data. This helps to learn useful latent representations of genomic DNA sequences that are then passed to the graph attention networks to be integrated over the adjacancy graphs derived from Hi-C/HiChIP contact matrices and to predict gene expression values (CAGE-seq). **C**. A 6Mb genomic region (11Mb-17Mb) of ch19 showing input and output signals and predictions in K562 cells, including epigenomic data (H3K4me3, H3K27ac, DNase), CAGE, HiChIP interaction graph, and predicted CAGE values for GraphReg and CNN models. Training and evaluations of the models are performed in the dashed middle 2Mb (here 13Mb-15Mb) region so that all genes can see the effects of their distal enhancers up to 2Mb.

For Seq-GraphReg (**Fig. 1B**, **Supplemental Fig. S1**), we feed one-hot-coded genomic DNA sequences to a series of 1D CNN, max-pooling, and ReLU layers, similar to Epi-GraphReg. The resulting bottleneck representations are then supplied to two blocks corresponding to different prediction tasks: a GAT block, similar to the Epi-GraphReg model, uses several GAT layers followed by a FC layer to predict the CAGE values; and a dilated CNN block, containing several dilated CNN layers whose dilated rate is multiplied by two each layer, to predict the 1D epigenomic data. Dilated CNNs have been used previously to increase the receptive field of CNN layers in deep learning models that predict epigenomic or expression data from DNA sequence, such as Basenji (Kelley et al., 2018), BPNet (Avsec et al., 2021b), ExPecto (Zhou et al., 2018), and Xpresso (Agarwal and Shendure, 2020). Therefore, Seq-GraphReg follows a multi-task learning approach to find more meaningful bottleneck representations to provide to the GAT block.

GATs have an advantage over other graph neural networks (GNN) in that they learn to weight edges in the graph from node features. In this way, GATs weigh enhancer-enhancer (E-E) and enhancer-promoter (E-P) interactions, based on the features learned in the promoter and enhancer bins, in order to predict CAGE values more accurately. However, non-attention-based GNNs such as graph convolutional networks (GCN) (Kipf and Welling, 2017; Bigness et al., 2021) fail to learn the importance of individual interactions. It has been shown that GATs outperform GCNs in other machine learning contexts as well (Veličković et al., 2018). We use different attention heads (**Fig. 1A,B**, **Methods**) in each GAT layer to enhance the flexibility of the model to learn distinct E-E and E-P interaction weights and to improve the prediction accuracy by integrating them together. **Fig. 1C** shows an example of a 6Mb region (11Mb-17Mb) of chromosome 19 in K562 cells with the corresponding true CAGE output, 1D epigenomic inputs, H3K27ac HiChIP interaction graph (FDR=0.1), and the predicted CAGE signals using Epi-GraphReg and Seq-GraphReg and their CNN counterparts, which we denote as Epi-CNN and Seq-CNN, respectively. The dashed middle 2Mb region in **Fig. 1C** indicates where we compute the loss in training and evaluate CAGE predictions. The inputs and outputs of Epi-CNN and Seq-CNN models are the same as those of Epi-GraphReg and Seq-GraphReg, respectively, with the exception that they use dilated CNN layers instead of GAT layers and do not use any 3D interaction data (**Supplemental Fig. S1**). We can train Seq-GraphReg in an end-to-end or separate manner. In separate training, instead of multi-task learning, we first predict epigenomic data from DNA sequence and then feed the bottleneck representation to the GAT layers to predict CAGE expression levels. Since CNN models to predict epigenomic signals can use smaller window sizes (100Kb instead of 6Mb), more filters can be used given the same GPU memory resources, yielding a better bottleneck representation, which consequently leads to improvement in CAGE prediction. However, this prediction improvement comes at the cost of losing access to fast backpropagation-based saliency scores at base-pair resolution (**Methods**).

### GraphReg accurately predicts gene expression by using 3D interaction data

We trained GraphReg models for three ENCODE human cell lines, GM12878, K562, hESC, and for mouse embryonic stem cells (mESC), for which complete 1D epigenomic data and 3D interaction data are available. To evaluate, we performed cross-validation experiments where we held out 2 chromosomes for testing, used 2 chromosomes for validation, and trained on all remaining autosomal chromosomes (**Methods**). Although our model predicts the CAGE signal at all genomic bins, we focused first on predictions for GENCODE-annotated TSS bins of protein coding genes, where the CAGE signal can be non-zero. **Figs. 2A,B** and **Supplemental Figs. S2-S8** show the CAGE prediction results for GM12878, K562, hESC, and mESC where predictions for test chromosomes across runs are concatenated together to obtain global performance results (20 test chromosomes over 10 runs). We computed the negative log-likelihood values (*NLL*), based on our loss function for the Poisson distribution, of the predicted CAGE signals for three gene sets: all genes (All); expressed genes, defined as CAGE signal ≥ 5 (Expressed); expressed genes with at least one E-P interaction (Interacting). We also reported Pearson correlation (*R*) of log-normalized predicted and true CAGE values (**Figs. 2A,B**, **Supplemental Figs. S2-S8**). We have also compared Seq-GraphReg and Seq-CNN with Basenji on K562 and GM12878 (**Supplemental Fig. S9**). In all cases, GraphReg models outperform the corresponding CNN models, achieving higher *R* and lower *NLL* (**Figs. 2A,B**, **Supplemental Figs. S2-S8**). After restricting to expressed or interacting genes, the problem gets harder and the prediction improvement of GraphReg over CNN models increases. Epigenome-based models (Epi-GraphReg and Epi-CNN) have higher prediction accuracy than the sequence-based models (Seq-GraphReg and Seq-CNN) (**Figs. 2A,B**, **Supplemental Figs. S2-S8**) since the epigenomic data are highly correlated with the CAGE output. To plot the boxplots in **Fig. 2** and assess statistical significance, we randomly sample 2000 predicted genes (in each category, All, Expressed, Interacting) 50 times, compute the mean of 2000 *NLL*s for both GraphReg and CNN, and perform a Wilcoxon signed-rank test to see if the *NLL*s are significantly smaller for Graphreg than CNN. We see that for both epigenome-based (**Fig. 2A**) and sequence-based (**Fig. 2B**) models, GraphReg has significantly smaller *NLL* than CNN (Wilcoxon signed-rank test), showing more accurate predictions for GraphReg.

**Figure 2:**
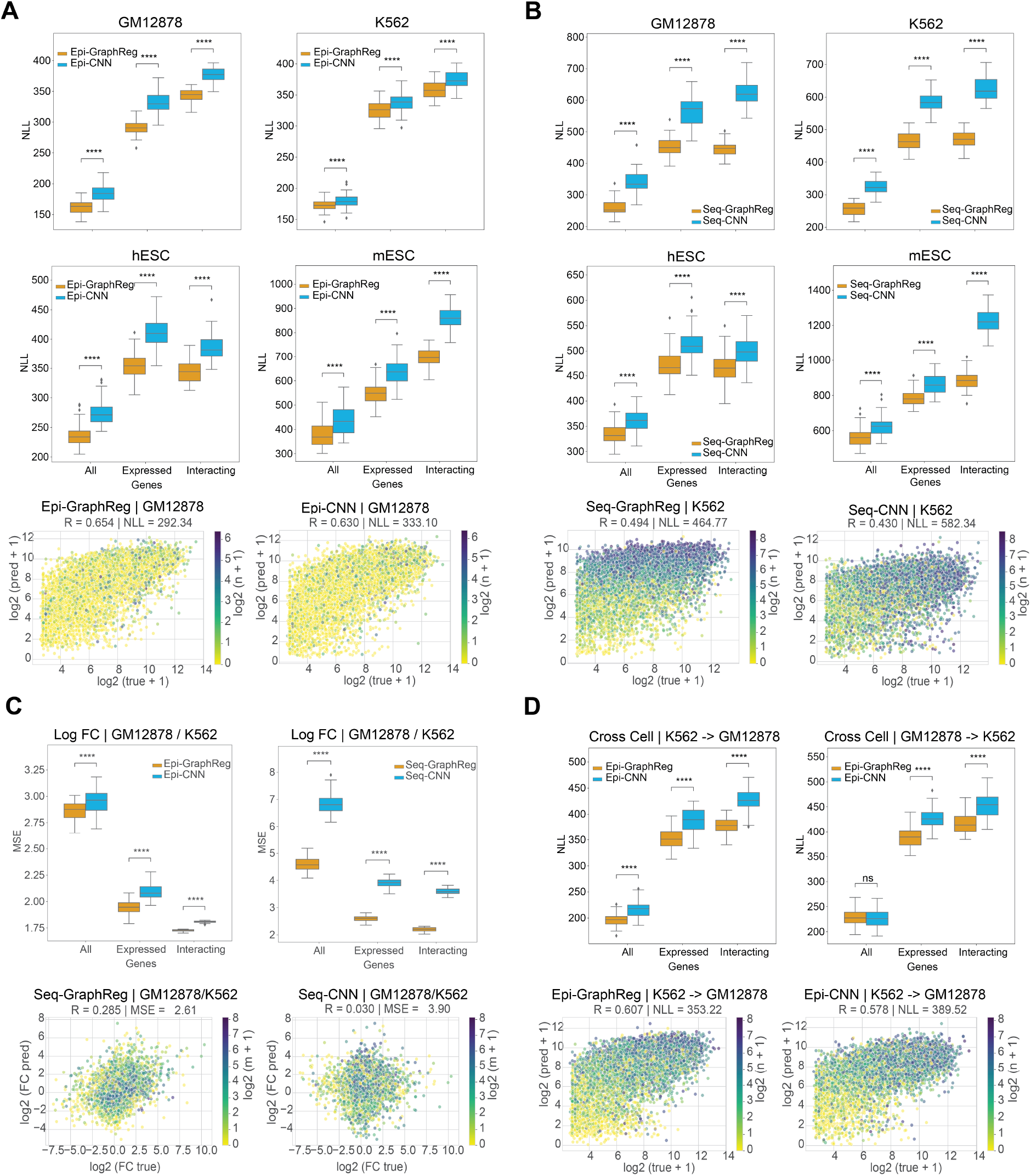
GraphReg models outperform their CNN counterparts for gene expression prediction. **A,B**. Negative log-likelihood (*NLL*, lower is better) between true and predicted CAGE signals of epigenome-based (**A**) and sequence-based (**B**) GraphReg and CNN models over 50 random selection of 2000 predicted genes from test chromosomes concatenated from 10 cross-validation experiments with different training, test, and validation chromosomes. Box plots show the distributions *NLL* in the cell lines GM12878, K562, hESC, and mESC for three gene sets: all genes, expressed genes (CAGE more than 5), and expressed genes with at least one 3D interaction (interacting). The 3D data used in Epi-GraphReg (**A**) for each cell type is as follows: Hi-C (FDR=0.001) for GM12878, HiChIP (FDR=0.01) for K562, Micro-C (FDR=0.1) for hESC, and HiChIP (FDR=0.1) for mESC. The 3D data used for Seq-GraphReg (**B**) is HiChIP (FDR=0.1) for GM12878, K562, and mESC, and is Micro-C (FDR=0.1) for hESC. Example scatter plots of all predicted test genes that are expressed (CAGE more than 5) are plotted for GM12878 in epigenome-based models (**A**) and K562 in sequence-based models (**B**). The sequenced-based models have been trained separately (and not using dilated CNN) for K562 and end-to-end for GM12878, hESC, and mESC. **C**. Box plots show mean squared error (*MSE*) of the true and predicted log-fold gene expression changes between GM12878 and K562 in 50 random selection of 2000 predicted genes from test chromosomes concatenated from 10 cross-validation experiments with different training, test, and validation chromosomes. The sets All, Expressed, and Interacting denote the intersections of such sets in GM12878 and K562. Both Epi- and Seq-GraphReg models have better prediction accuracy than their CNN counterparts. The scatter plots of the true log-fold gene expression changes and the log-fold changes derived from the predicted CAGE values by Seq-GraphReg and Seq-CNN, between GM12878 and K562, are shown for expressed genes (CAGE more than 5 in both K562 and GM12878). TSS bins are color-coded by the minimum number of 3D interactions in GM12878 and K562 (*m*). Seq-GraphReg has higher *R* and lower *MSE* than Seq-CNN. **D**. Epi-GraphReg models demonstrate higher cell-to-cell generalization capability than Epi-CNN models. Box plots show the distributions of *NLL* on the test cell type (K562 or GM12878) when trained on the other cell type over 50 random selection of 2000 predicted genes from test chromosomes of the test cell concatenated from 10 cross-validation experiments with different training, test, and validation chromosomes in the training cell. The models are evaluated on the same test chromosomes in the unseen test cell. HiChIP (FDR=0.1) is used for both cells. The generalization of Epi-GraphReg from K562 to GM12878 is significantly better (*p* < 10^−4^, Wilcoxon signed-rank test) than Epi-CNN in all gene sets. The generalization of Epi-GraphReg from GM12878 to K562 is significantly better (*p* < 10^−4^, Wilcoxon signed-rank test) than Epi-CNN in expressed and interacting genes. The scatter plots of all predicted test genes that are expressed (CAGE more than 5) are plotted when trained on K562 and tested on GM12878.

We also examined scatter plots of log-normalized (log_2_(*x* + 1)) predicted versus true CAGE values of GraphReg and CNN models across the expressed genes in mESC, hESC, K562, and GM12878 (**Figs. 2A,B**, **Supplemental Figs. S3-S8**). Here TSS bins are color-coded by their number of 3D interactions (*n*). An example of Epi-GraphReg and Epi-CNN on GM12878 using Hi-C (FDR=0.001) is shown in **Fig. 2A**, and an example of Seq-GraphReg and Seq-CNN on K562 using HiChIP (FDR=0.1) with separate training and no dilated layers is shown in **Fig. 2B**. For epigenome-based models, in **Fig. 2A**, Epi-GraphReg yields *R* = 0.654 and *NLL* = 292.34 on expressed genes, outperforming Epi-CNN with *R* = 0.63 and *NLL* = 333.10. For sequence-based models, in **Fig. 2B**, Seq-GraphReg yields *R* = 0.494 and *NLL* = 464.77 on expressed genes, outperforming Seq-CNN with *R* = 0.43 and *NLL* = 582.34. More scatter plots for different GraphReg models can be found in **Supplemental Figs. S3-S8**. Analogous to our previous notion of *gene regulatory complexity* (González et al., 2015), we refer to genes with *n* = 0 as “simple genes” and those with *n* > 0 (at least one 3D interaction) as “complex genes”. We hypothesized that by exploiting 3D interaction data, GraphReg would better model the regulation of complex genes. This fact can be easily seen in scatter plots shown in **Fig. 2B**.

Notably, the extent of improvement of Seq-GraphReg over Seq-CNN is usually higher than that of the Epi-GraphReg over Epi-CNN (**Figs. 2A,B**, **Supplemental Figs. S2-S8**), suggesting the importance of 3D interaction data for integrating distal regulatory elements and improving prediction accuracy when predicting from DNA sequence, especially for more complex genes with many E-P interactions (**Supplemental Figs. S7-S8**). In particular, Seq-CNN models predict lower values for some highly expressed genes with many 3D interactions, while Seq-GraphReg can more accurately predict these higher expression values (**Fig. 2B**, **Supplemental Figs. S7-S8**). The effects of 3D assay (Hi-C, HiChIP, Micro-C) and different FDR levels (0.1, 0.01, 0.001, related to various noise levels in the graphs) for Epi-GraphReg is shown in **Supplemental Figs. S2-S6**.

GraphReg’s improved performance over CNNs can also be witnessed by calculating the log-fold change between CAGE predictions for two different cell types (**Fig. 2C**), and by comparing to true log-fold changes using mean squared error (*MSE*). Evaluation of predicted log-fold changes in GM12878 versus K562 on held-out chromosomes in 10 cross-validation runs (**Fig. 2C**) confirmed the improvement of Epi-GraphReg and Seq-GraphReg over Epi-CNN and Seq-CNN, respectively. We see in **Fig. 2C** that *MSE*s of GraphReg are significantly smaller than CNN (Wilcoxon signed-rank test), and the performance improvements are higher for Seq-GraphReg versus Seq-CNN. **Fig. 2C** shows the scatter plots of predicted versus true log-fold change in TSS bins on held-out chromosomes for expressed genes (CAGE ≥ 5 in both GM12878 and K562), color-coded by the minimum number of 3D interactions over the two cell types (*m*). For Seq-GraphReg, HiChIP (FDR=0.1) is used in both GM12878 and K562. For Epi-GraphReg, Hi-C (FDR=0.001) and HiChIP (FDR=0.01) are used in GM12878 and K562, respectively. As shown in scatter plots of **Fig. 2C**, Seq-GraphReg achieves *R* = 0.285 and *MSE* = 2.61 in expressed genes, compared to very poor *R* = 0.03 and *MSE* = 3.9 for Seq-CNN. Scatter plots of true versus predicted log-fold change by Epi-GraphReg and Epi-CNN are shown in **Supplemental Fig. S10**, with *R* = 0.561 and *MSE* = 1.94 for Epi-GraphReg and *R* = 0.535 and *MSE* = 2.08 for Epi-CNN. Overall, these results confirm that using 3D chromatin interactions in GraphReg leads to improved gene expression prediction.

We investigated how well the cell-type agnostic Epi-GraphReg and Epi-CNN models can generalize from one cell type to another. **Fig. 2D** shows the box plots of loss (*NLL*) when we train on one cell type (K562 or GM12878) and test on the other, using 10 cross-validation runs with distinct sets of validation chromosomes (held out from training and used to assess performance in the test cell type). Epi-GraphReg significantly outperforms Epi-CNN (*p* < 10^−4^, Wilcoxon signed-rank test) for generalization from K562 to GM12878 and vice versa in all categories except one: all genes from GM12878 to K562. Example scatter plots for these cross-cell-type and cross-chromosome generalization tasks show the improvement of Epi-GraphReg over Epi-CNN (**Fig. 2D**): when training on K562 with HiChIP (FDR=0.1) and testing on GM12878 with HiChIP (FDR=0.1), Epi-GraphReg attains *R* = 0.607 and *NLL* = 353.22, whereas Epi-CNN yields *R* = 0.578 and *NLL* = 389.52 for expressed genes. The effect of 3D assay choice and FDR values for cross-cell-type generalization is examined in **Supplemental Figs. S11-S13**, where we see that the best and most robust generalization happens when we use HiChIP in both train and test cell types or HiChIP in train and Hi-C in test cell types, and the worst and least robust generalization performance occurs when we use Hi-C in train and HiChIP in test cell types. One reason for this phenomenon could be that HiChIP interactions, which are enriched for enhancer-promoter interactions, are more generalizable between cell types than Hi-C interactions, which depend strongly on the depth of coverage and library complexity and may be dominated by structural interactions in lower coverage data sets. Similarly to K562 → GM12878, when training on GM12878 with HiChIP (FDR=0.001) and testing on K562 with HiChIP (FDR=0.1), Epi-GraphReg attains *R* = 0.563 and *NLL* = 389.71 compared to *R* = 0.585 and *NLL* = 425.91 for Epi-CNN on expressed genes (**Supplemental Fig. S13**).

We also wanted to determine the classes of genes for which GraphReg’s predictions are better than those of CNN models and thus understand the genes for which 3D information gives the most benefit. To this end, we defined a simple metric called *Delta_NLL_* = *NLL_CNN_* − *NLL_GraphReg_*. If *Delta_NLL_* > 0 for a gene, it means that GraphReg’s prediction is better than CNN for that gene, and if *DeltaNLL* < 0 for a gene, it means that CNN’s prediction is better than GraphReg for that gene. We produced scatter plots of *DeltaNLL* versus the number of interactions for all expressed genes in all four cell types GM12878, K562, hESC, and mESC, and for both sequence-based (Seq-GraphReg and Seq-CNN) and epigenome-based (Epi-GraphReg and Epi-CNN) models (**Supplemental Figs. S3-S8 and Figs. S12-S13**). We can see that the majority of genes have positive *DeltaNLL*, especially the ones with more interactions. Furthermore, the number of genes with positive *DeltaNLL* is higher in sequenced-based models, showing the greater advantage of using 3D information and GraphReg when predicting from DNA sequence. We have also listed in **Supplemental Figs. S14-S21** the top genes for which either GraphReg or CNN predicts better.

### GraphReg accurately identifies functional enhancers of genes

Thanks to the feature attribution methods for machine learning models – e.g. saliency maps (gradient-by-input), DeepLIFT (Shrikumar et al., 2017), DeepSHAP (Lundberg and Lee, 2017), and Integrated Gradients (Sundararajan et al., 2017) – it is possible to derive the important input features for the prediction of a specific output. Since GraphReg allows each gene to be influenced by its potential enhancers through interactions in the 3D graph, we hypothesized that feature attribution analysis would allow the identification of functional distal enhancers, i.e. those that contribute to the regulation of target genes. CRISPRi-FlowFISH (Fulco et al., 2019) is a recent enhancer screening approach that uses KRAB-dCas9 interference with candidate enhancer regions in pooled fashion and RNA FISH against a gene of interest as readout; this enables estimation of the effect size of perturbing each candidate enhancer on target gene expression. The developers of CRISPRi-FlowFISH introduced a score called Activity-by-Contact (ABC) for finding and ranking enhancers of a gene, which is considered the current state-of-the-art for this problem. The ABC score is the product of “activity”, defined as the geometric mean of DNase and H3K27ac signal in each candidate enhancer region, and “contact”, defined by the KR-normalized Hi-C signal between the target promoter and the candidate enhancer, normalized so that the ABC scores of all candidate enhancers up to 5Mb from the gene sum to one.

To determine functional enhancers for genes, we used two feature attribution methods: the saliency map (gradient-by-input) and DeepSHAP (Lundberg and Lee, 2017). For the epigenome-based models (Epi-GraphReg and Epi-CNN), as the inputs are binned at 100bp resolution, both saliency and DeepSHAP scores are computed at 100bp resolution as well. We used all-zero data as the reference signal for DeepSHAP so that it can evaluate the importance of peak regions in the input tracks. For the sequence-based models (Seq-GraphReg and Seq-CNN), the feature attribution methods can provide scores at nucleotide resolution. However, for enhancer scoring and validation with FlowFISH analysis, as suggested in Basenji (Kelley et al., 2018), we derived the saliency maps as the dot product of the 100bp bin representations (bottleneck) and the gradient of the model prediction for the gene with respect to those bin representations. **Fig. 3A** shows the distribution of area under the precision-recall curve (auPR) values for 19 genes from the K562 FlowFISH dataset with more than 10 candidate enhancers for all models. The overall precision-recall curve across 2574 enhancer-gene (E-G) pairs for all 19 genes is shown in **Fig. 3B**. HiChIP (FDR=0.1) has been used for GraphReg in these experiments. These results confirm that feature attribution applied to GraphReg models more accurately identifies the functional enhancers of the genes from a pool of candidate enhancers than the ABC score or feature attribution on the corresponding CNN models (**Fig. 3A,B**). The highest auPR among GraphReg models was 0.4236 (Epi-GraphReg with saliency), outperforming the best CNN model (auPR 0.366 for Epi-CNN with saliency) and ABC (auPR 0.364).

**Figure 3:**
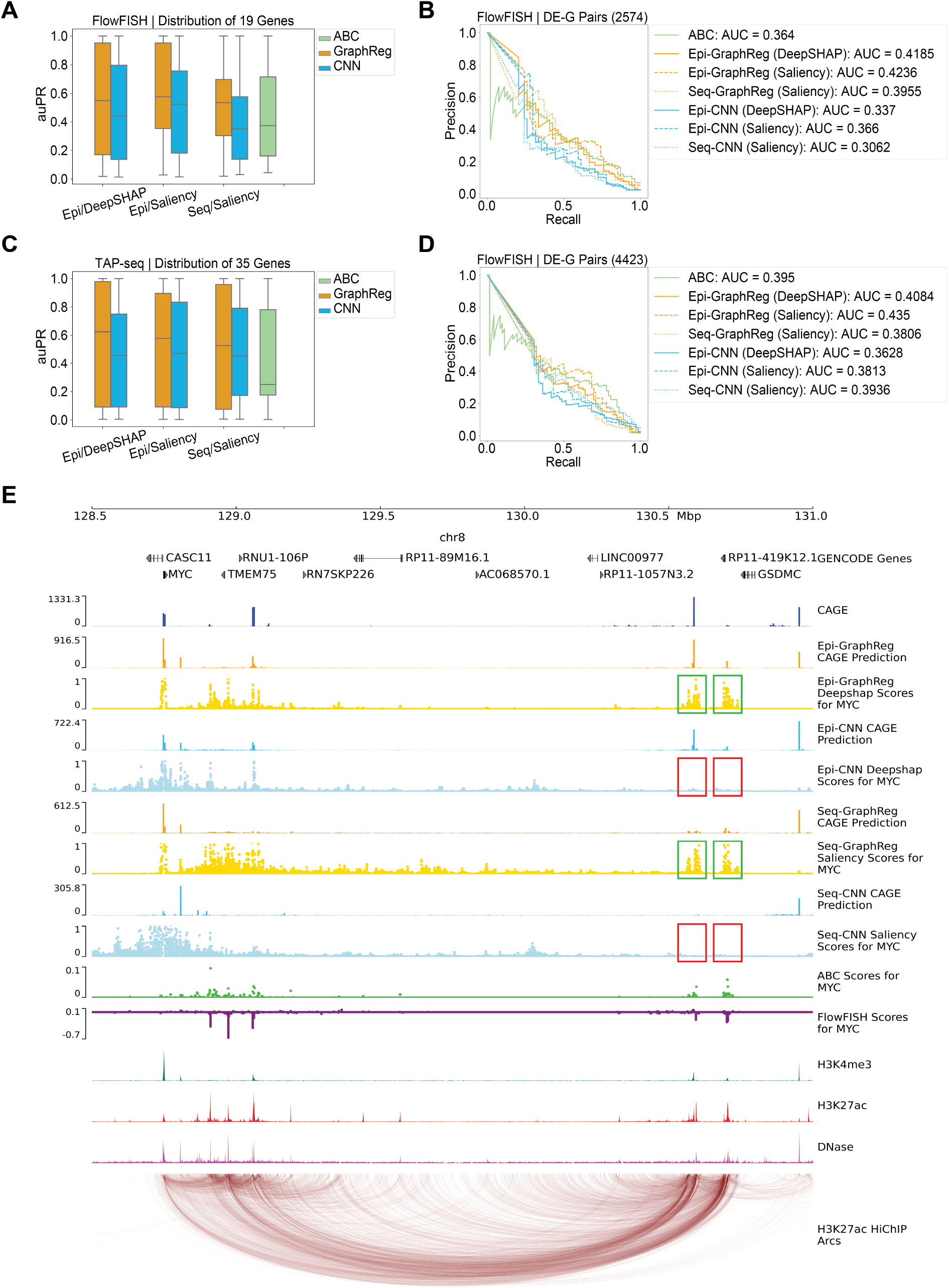
GraphReg models more accurately identify functional enhancers of genes. **A**. Distribution of the area under the precision-recall curve (auPR) for 19 genes in K562 cells based on CRISPRi-FlowFISH data. GraphReg models have higher median of auPR than both CNN and Activity-by-Contact (ABC) models. **B**. Precision-recall curves of the GraphReg, CNN, and ABC models for identifying enhancers of 19 genes screened by CRISPRi-FlowFISH. **C**. Distribution of auPR for 35 genes in K562 cells based on TAP-seq data. GraphReg models have higher median of auPR than both CNN and ABC models. **D**. Precision-recall curves for the GraphReg, CNN, and ABC models for identifying functional enhancers of 35 genes as determined by TAP-seq. **E**. *MYC* locus (2.5Mb) on chr8 with epigenomic data, true CAGE, predicted CAGE using GraphReg and CNN models, HiChIP interaction graph, and the saliency maps of the GraphReg and CNN models, all in K562 cells. Experimental CRISPRi-FlowFISH results and ABC values are also shown for *MYC*. Feature attribution shows that GraphReg models exploit HiChIP interaction graphs to find the distal enhancers, while CNN models find only promoter-proximal enhancers. Green and red boxes show true positives and false negatives, respectively. CNN models miss the distal enhancers and consequently lead to false negatives in very distal regions.

To assess the ability of models to find distal enhancers, we examined the example of *MYC* and considered only candidate enhancers more than 10Kb from TSS. There are 200 such candidate distal enhancer-gene (DE-G) pairs for *MYC*, including 8 true functional enhancers. Precision-recall analysis for *MYC* (**Supplemental Fig. S22A**) found the highest auPR of GraphReg models to be 0.576 (Epi-GraphReg with DeepSHAP), strongly outperforming the best CNN model (auPR 0.0792, Epi-CNN with DeepSHAP) as well as ABC (auPR 0.3567). Visualization of the feature attribution scores gives insight into how 3D information allows GraphReg to access distal regulatory information that dilated CNNs cannot exploit, despite their large receptive field. **Fig. 3E** shows a 2.5Mb genomic region containing the *MYC* locus, with the corresponding epigenomic data, HiChIP graph edges, and true and predicted CAGE signals by GraphReg and CNN models in K562 cells, together with DeepSHAP and saliency scores of the four models for *MYC*, and ABC scores and FlowFISH results for *MYC*. Large negative scores in the FlowFISH track indicate true functional enhancers for *MYC*. Feature attribution tracks in **Fig. 3E** show that GraphReg models are able to capture the most distal enhancers of *MYC* (green boxes), about 2Mb away from its TSS, while the CNN models fail to capture these enhancers and produce false negatives (red boxes).

We further evaluated GraphReg using recent chromosome-wide enhancer screening data from TAP-seq (targeted perturb-seq) for chromosomes 8 and 11 in K562 cells (Schraivogel et al., 2020). We restricted to the 35 genes with at least one functional enhancer, considering all screened enhancers whose distance from the target gene TSS is less than 2Mb. **Fig. 3C** shows the distribution of auPR for these 35 genes. Similar to CRISPRi-FlowFISH results (**Fig. 3A**), GraphReg models have better median auPR than ABC scores or the CNN models using either DeepSHAP or saliency as the feature attribution method. Pooling all the E-G pairs of these 35 genes (4423 E-G pairs in total), precision-recall analysis (**Fig. 3D**) shows that the highest auPR for the GraphReg models is 0.435 (Epi-GraphReg with saliency), outperforming the best CNN model (auPR 0.3936) and ABC (auPR 0.395).

As another example of distal enhancer discovery in the TAP-seq dataset, we considered screened enhancers more than 20Kb from TSS of *IFITM1* (79 distal DE-G pairs, 5 true functional enhancers). Precision-recall analysis for *IFITM1* shows that the highest auPR among the GraphReg models is 0.894 (Epi-GraphReg with saliency), compared to 0.635 for the best CNN model (Epi-CNN with DeepSHAP) and 0.751 for ABC (**Supplemental Fig. S22B**). We also plotted a 250Kb genomic region containing the *IFITM1* locus, with the corresponding K562 epigenomic data, HiChIP graph edges, true and predicted CAGE signals by GraphReg and CNN models, feature attribution scores for *IFITM1*, ABC scores for *IFITM1*, and TAP-seq results for *IFITM1* (**Supplemental Fig. S22C**). Large negative scores in the TAP-seq track indicate true functional enhancers of *IFITM1*. Here all GraphReg, CNN, and ABC models are able to capture the distal enhancers of *IFITM1* (green boxes), but the CNN models also capture many non-functional enhancers as false positives (red boxes).

### Seq-GraphReg predicts the direct targets of transcription factor perturbation via motif ablation

To assess whether Seq-GraphReg captures meaningful transcription factor (TF) motif information, we asked whether we could predict differential gene expression in TF perturbation experiments by applying our model to genomic sequences where the TF’s binding motif had been ablated. We downloaded RNA-seq data for 51 CRISPRi TF knockout (KO) experiments in K562 cells from ENCODE and retained 29 experiments for which at least 200 genes were significantly downregulated. Using two trained models each for Seq-GraphReg and Seq-CNN, we predicted gene expression using wildtype genomic sequences and sequences where hits of a given TF motif had been zeroed out, then performed differential expression analysis on these values to predict the 100 most downregulated target genes for the corresponding TF KO experiment (**Methods**). To have confident true labels, we restricted this analysis to genes that have true significant differential expression (up- or downregulated) in the real respective TF KO experiments and that also have wildtype CAGE value of at least 20.

**Fig. 4A** shows the distributions of true mean logFC of the 100 predicted target genes over all TFs for the Seq-GraphReg and Seq-CNN models. As a baseline, we report the distribution of true mean logFC over all significantly differential genes (*p_adj_* < 0.05). This baseline actually shows the average performance of a random algorithm that chooses 100 genes randomly out of all significantly differential genes. We also show the corresponding distributions after restricting to predicted genes with *n* ≥ 5, where *n* denotes the number of 3D interactions. In both settings, Seq-GraphReg’s predicted target genes are significantly more downregulated than those of Seq-CNN’s or than baseline performance (Wilcoxon signed-rank test on distributions of mean logFC values). **Fig. 4B** shows the heatmaps of the true mean logFC of the top 100 predicted genes by Seq-GraphReg and Seq-CNN for each TF, for *n* ≥ 0 and *n* ≥ 5. For the majority of TFs, the mean logFC of Seq-GraphReg’s predicted targets is more negative than that of Seq-CNN’s predicted targets, validating the improved performance of Seq-GraphReg for identifying true downregulated genes. **Fig. 4C** shows the distributions of the precision (fraction of true significantly downregulated genes among the 100 predicted genes) for all TFs for Seq-GraphReg and Seq-CNN, for *n* ≥ 0 and *n* ≥ 5. As baseline, we report the fraction of significantly downregulated genes (*p_adj_* < 0.05 and logFC <0) over all significantly differential genes (*p_adj_* < 0.05), for each TF. This baseline shows the average performance of a random algorithm. Again, the precision values are always highest for Seq-GraphReg and significantly greater than for Seq-CNN and baseline (Wilcoxon signed-rank test). **Fig. 4D** shows this data in heatmap form, confirming that for the majority of TFs, the precision for Seq-GraphReg is higher than for Seq-CNN.

**Figure 4:**
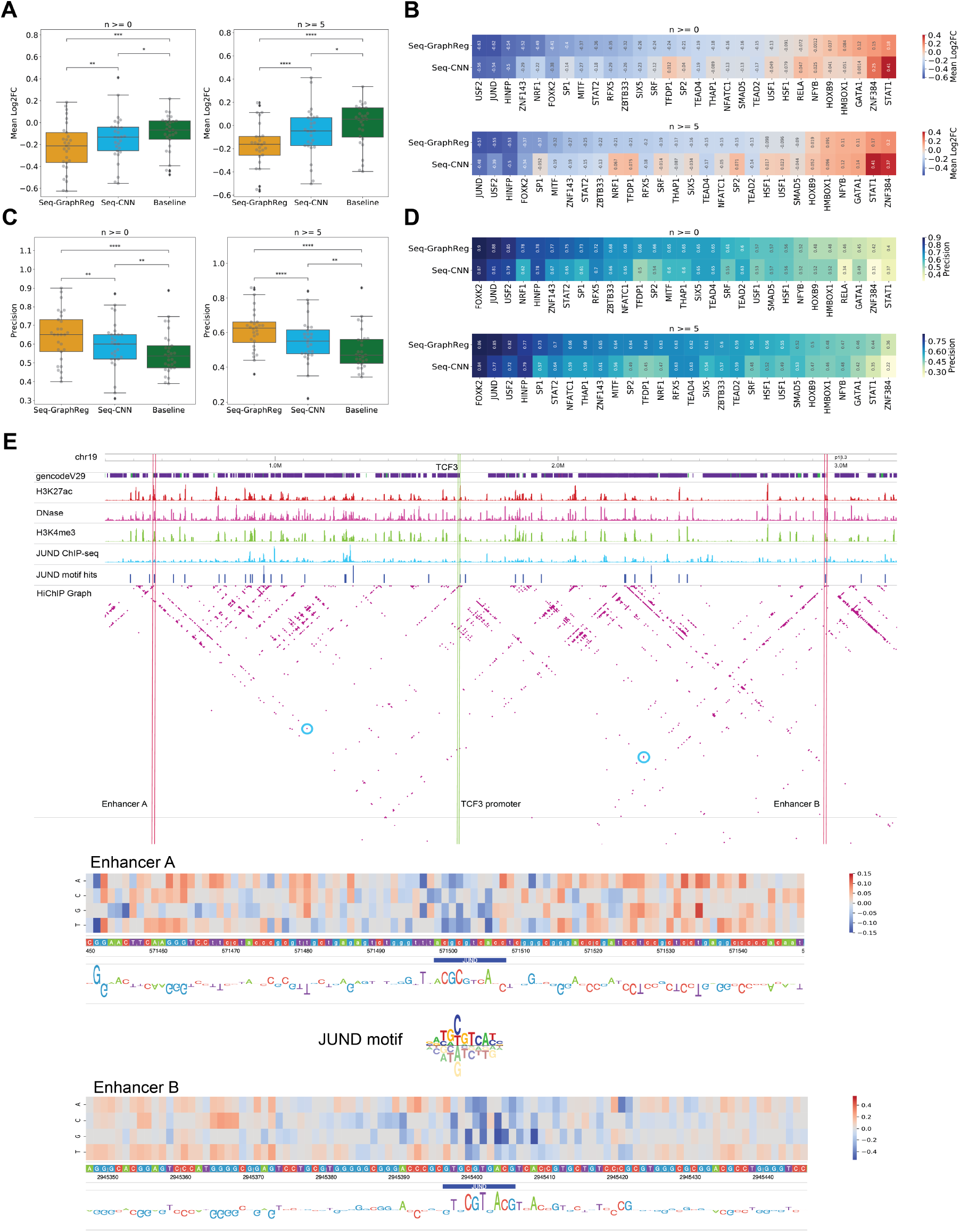
Seq-GraphReg accurately predicts the regulatory effects of transcription factor knockouts by in silico motif ablation. **A**. Distributions of true mean logFC (over 100 predicted target genes) over all TFs for Seq-GraphReg, Seq-CNN, and baseline for *n* ≥ 0 and *n* ≥ 5, where *n* denotes the number of enhancer-promoter (E-P) interactions. The median of true mean logFCs for predicted genes by Seq-GraphReg is negative, and the distribution is significantly more downregulated than that of Seq-CNN and baseline (Wilcoxon signed-rank test). **B**. Heatmaps of the true mean logFC of top 100 predicted genes by Seq-GraphReg and Seq-CNN for each TF, for *n* ≥ 0 and *n* ≥ 5. For the majority of TFs, the mean logFC of Seq-GraphReg’s predicted targets is more negative than that of Seq-CNN’s targets. **C**. Distributions of precision values (fraction of true significantly down-regulated genes among 100 predicted genes) of all TFs for Seq-GraphReg, Seq-CNN, and baseline, for *n* ≥ 0 and *n* ≥ 5. The precision is always highest in Seq-GraphReg and significantly greater than for Seq-CNN and baseline (Wilcoxon signed-rank test). **D**. Heatmaps of precision values (fraction of true significantly down-regulated genes among top 100 predicted genes) for Seq-GraphReg and Seq-CNN for each TF and for *n* ≥ 0 and *n* ≥ 5. For the majority of TFs, the precision of Seq-GraphReg is higher than Seq-CNN. **E**. A visual example of the effect of JUND KO on the gene *TCF3*. JUND motif hits around *TCF3* are plotted in blue bars. The promoter of *TCF3* is indicated by green lines, and two distal enhancers A (1.08 Mb downstream) and B (1.29 Mb upstream) of the gene *TCF3* by red lines. The interactions of enhancers A and B with the promoter of *TCF3* in HiChIP graph are marked by blue circles. In silico mutagenesis (ISM) is performed in 100 bp regions of enhancers A and B, each centered at a JUND motif, and the ISM heatmaps are shown. The heatmaps show the difference in predictions (mutated - reference) after applying a mutation at each nucleotide. The heatmaps around the JUND motif in both enhancers A and B are blue-ish, meaning that ISM reflects the importance of JUND motif in these regions for *TCF3* expression prediction. The base-level representations of ISM scores are the negative of summation of all 4 scores (only 3 are non-zero) at each nucleotide, which identify the JUND motifs.

**Fig. 4E** depicts a visual example of predicting the effect of JUND KO on the gene *TCF3* in K562. *TCF3* is among top 20 predicted genes of Seq-GraphReg for the JUND TF (**Supplemental Fig. S23A**), and CRISPRi KO of JUND leads to significant downregulation of *TCF3*. **Fig. 4E** shows JUND motif hits (blue bars) around *TCF3*, indicating the motifs ablated for in silico prediction, along with JUND ChIP-seq data in K562 cells, demonstrating that the motif hits indeed fall in JUND ChIP-seq peaks. In order to see if Seq-GraphReg can identify the JUND motifs in distal enhancers we did the following. First, we chose two distal enhancers (indicated in **Fig. 4E** by A and B) of the gene *TCF3* with direct (one-hop) enhancer-promoter interactions in the HiChIP graph. Enhancers with direct interactions will have a bigger effect size on gene expression upon mutation. We also visualize graph heatmaps (adjacency matrices) instead of loops to facilitate viewing the interactions. Second, we used in silico mutagenesis (ISM) for feature attribution. We performed ISM on Seq-GraphReg trained with FFT (fast Fourier transform) loss to get more meaningful and interpretable motifs. We have borrowed the idea of adding FFT loss from a recent study (Tseng et al., 2020). We used FFT in the separate training scheme of Seq-GraphReg, instead of end-to-end training, due to GPU memory limitation. **Fig. 4E** shows in silico mutagenesis results for the two distal enhancer regions A (1.08 Mb downstream) and B (1.29 Mb upstream) of the gene *TCF3*, whose interactions with the *TCF3* promoter are marked by blue circles in the HiChIP graph. The promoter of *TCF3* is specified by a green line, and the enhancers A and B are specified by red lines. ISM is performed in 100 bp regions of enhancers A and B, each centered at a JUND motif. The ISM heatmaps show the difference in predictions (mutated - reference) after applying a mutation at each nucleotide. The heatmaps around the JUND motif in both enhancers A and B are blue-ish, meaning that ISM identifies the importance of the JUND motif in these regions for target gene expression prediction. In other words, mutations in the JUND motif in distal enhancers A and B lead to reduction in expression prediction, showing the positive effect of these distal JUND motifs on the expression of *TCF3*. The base-level representations of ISM scores are the negative of summation of all 4 scores (only 3 are non-zero) at each nucleotide, which identify the JUND motifs.

Overall, these results show that Seq-GraphReg models can better capture the regulatory relationships of TFs and genes than Seq-CNN models. This validates the primary motivation for our model, namely that TFs can regulate their target genes by binding distal enhancers that loop to target promoters, and that these regulatory effects cannot be effectively learned without 3D interaction data. GraphReg is therefore the model of choice to capture complex gene regulatory relationships.

## Discussion

Until now, machine learning models for the genome-wide prediction of gene expression in a given cell state, or gene expression changes in a cell state transition, have largely relied on 1D epigenomic data and/or genomic DNA sequence without using 3D genomic architecture. Prior models include regularized linear regression approaches (González et al., 2015; Osmanbeyoglu et al., 2019) as well as more recent work using convolutional neural networks (Avsec et al., 2021a; Kelley et al., 2018; Singh et al., 2016; Zhou et al., 2018; Agarwal and Shendure, 2020). However, these previous linear models use a fixed assignment of regulatory elements to genes without incorporating 3D interaction data, while current deep learning models consider relatively local features, such as promoters and at most nearby enhancers, and therefore cannot capture the impact of distal regulatory elements, which can be 1Mb or farther away from gene promoters. There are two studies that tried to use 3D data to predict gene expression (Bigness et al., 2021; Zeng et al., 2019), but they failed to address important aspects of modeling gene regulation. The most directly relevant one, GC-MERGE (Bigness et al., 2021), uses histone modification and Hi-C data to predict gene expression (RNA-seq) using graph convolutional networks (GCN); however, they do not provide any insight about gene regulation rules, such as finding functional enhancers or revealing the role of TF binding motifs on gene regulation. Our proposed method, GraphReg, is the most comprehensive and versatile model that is capable of answering interesting biological questions regarding gene regulation. We have provided the properties of previous deep learning models for gene expression prediction in **Supplemental Table S1**. The comparisons of GraphReg with these methods in terms of Pearson correlation (R) of predicted versus true gene expression values are provided in **Supplemental Tables S1-S2** and **Supplemental Fig. S9**.

We have shown that GraphReg more effectively models the impact of distal enhancers on gene expression than 1D dilated CNNs. While dilated CNNs have a large receptive field, our feature attribution analyses show that only relatively promoter-proximal information influences gene expression prediction in these models. By contrast, GraphReg exploits 3D chromatin interactions to access distal information up to 2Mb from the gene promoter. In other words, GraphReg adds inductive bias from biology to the deep predictive model, which converts a black-box model attending to everywhere to a more focused model attending to relevant regions.

An alternative attention-based methodology for learning long-range interactions in sequential input is the tranformer, a model that provides state-of-the-art performance in machine translation and other natural language processing tasks (Vaswani et al., 2017). A recent work, Enformer (Avsec et al., 2021a), introduced a transformer model for the prediction of CAGE as well as epigenomic tracks from genomic sequence. Currently, however, this transformer-based architecture can integrate information up to 100Kb away in the genome, an order of magnitude less than GraphReg, while requiring considerable computational resources to train. GraphReg’s use of 3D interaction data through graph attention networks therefore provides an efficient and biologically well-motivated means to encode distal regulation.

We showed that Seq-GraphReg meaningfully encodes TF motif binding information by performing motif ablation experiments, where predicted differential gene expression based on ablated versus wildtype genomic sequences recovered true regulatory effects as measured by CRISPRi TF knockout experiments. A longer term goal of this work is to model the impact of distal regulatory variants on gene expression, extending efforts to predict the regulatory impact of promoter-proximal genetic variants (Zhou et al., 2018). We anticipate that it will be critical to train such models using true genetic sequence variation and expression in disease-relevant cellular contexts as these data become available.

While we currently use GraphReg to predict bulk CAGE-seq counts with Poisson loss, other transcriptomic assays could be explored in future work. For example, suitably processed nascent transcription assays such as GRO-seq would yield a quantification of promoter activity as well as enhancer RNA expression and could potentially be used to train GraphReg models. We also envision extensions to single cell multi-omic data sets – training at the pseudo-bulk level where signals are aggregated over cell clusters or meta-cells, or even at the single-cell level. Here (aggregated) scRNA-seq gives a tag-based output signal similar to CAGE-seq. Therefore, we anticipate that GraphReg models will have broad applicability for interpreting the function of epigenomic and genomic variation on gene expression.

## Methods

### CNN layers for learning local representations of 1D data

The inputs to the first layer of the Epi-GraphReg (Epi-CNN) and Seq-GraphReg (Seq-CNN) models are 1D epigenomic data and genomic DNA sequence, respectively. Regardless of the 1D input type, we use several CNN layers followed by RELU activation, BatchNorm, dropout, and max-pooling to learn local representations for 5Kb bins of the genome (see **Supplemental Fig. S1**). We consider genomic regions of 6Mb in length as our input; hence, we have vectors of size *N* = 1200, representing 5Kb bins over the 6Mb region, for CAGE values. We bin the epigenomic data at 100bp resolution; therefore, the length of each of epigenomic data would be 60000. In the GraphReg models, we define the set of local representations by *H* = {*h*_1_, *h*_2_, …, *h_N_*}, where *h_i_* ∈ ℝ^*F*^ and *F* = 128 is the channel number of the last CNN layer just before the GAT block. Then *H* is given to the GAT block, which is a type of graph neural network (GNN) (Kipf and Welling, 2017; Xu et al., 2019; Veličković et al., 2018), where *h_i_* is the node feature of node *i*. To make the comparisons fair, in Epi-CNN and Seq-CNN models we use 8 layers of dilated CNNs with residual connections (Kelley et al., 2018; Avsec et al., 2021b) to increase the receptive fields up to 2.5Mb upstream and downstream of gene TSSs, where the dilation rate is multiplied by 2 at each layer. In the sequence-based models (Seq-GraphReg and Seq-CNN) we also predict epigenomic data in order to guide the models to learn informative local motif representations; after first 5 CNN layers, where the resolution is 100bp, we use 6 layers of dilated CNNs with residual connections to predict three epigenomic assays, H3K4me3, H3K27ac, and DNase at 100bp resolution. (see **Supplemental Fig. S1**). The learned local features just before 6 dilated CNN layers are called bottleneck representations that are passed through some CNN layers with max-pooling to bring the resolution to 5Kb and then are given to GAT block in Seq-GraphReg (and 8 layers of dilated CNN in Seq-CNN) to predict the CAGE (see **Supplemental Fig. S1**).

### Graph attention networks for integration of local features via 3D interaction data

We employ the graphs extracted from 3D data (Hi-C, HiChIP, Micro-C) in order to capture gene regulatory interactions and predict gene expression (CAGE-seq). We processed 3D data sets at 5Kb resolution using HiC-DC+ (Carty et al., 2017; Sahin et al., 2021) and kept only the significant interactions with three significance levels of FDR less than 0.1, 0.01, and 0.001. The nodes in the extracted graphs therefore represent 5Kb genomic bins and the edges indicate the most significant interactions between bins. Each 6Mb genomic input region thus corresponds to a graph with *N* = 1200 nodes whose features *H* = {*h*_1_, *h*_2_, …, *h_N_*} have been learned in the previous CNN block.

The graph attention network (GAT) block receives a graph *G* = (*V, E*) and a set of node features 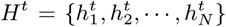 from the previous layer *t* and outputs an updated set of node features 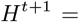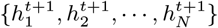. Each GAT layer uses two weight matrices: 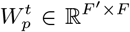 for promoters (or self nodes) and 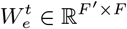 for enhancers (or neighbor nodes), where *F*′ is the number of output features in each GAT layer. We know that the importance of distinct enhancers on a promoter might be different. In the case of GCNs, we treat all the enhancers as of equal importance because all the E-P weights are the same. However, by using GAT, we give the model the ability to adjust the weights of different enhancers based on their importance for the task of predicting expression of a specific gene. Therefore, the GAT formulation makes more biological sense for our problem. Note that here, unlike previous graph neural networks (Veličković et al., 2018), we do not include self-loops in the graph *G*. We have decoupled self-loops and neighbor-loops due to the differing role of promoters and enhancers in the model. By only including the neighbor-loops in the graph *G*, the model focuses on distal enhancers that cannot be captured in local models such as CNNs.

We define the self-attention mechanism from the nodes *j* to the node *i* at layer *t* as

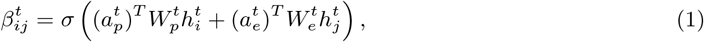

where 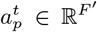 and 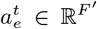 are two weight vectors, *σ*(.) is the sigmoid function, 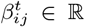 is the attention weight from node *j* to node *i* at layer *t*, and *T* denotes the transpose of a vector. By using a sigmoid function instead of a conventional softmax function, we give extra freedom to the model to discard irrelevant and non-enhancer interactions. We also account for the cardinality of the nodes by defining 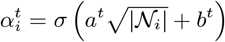, where *a^t^* ∈ ℝ^*F′*^ and *b^t^* ∈ ℝ^*F′*^ are two weight vectors. Finally, we define the updates of the node features at the next layer as

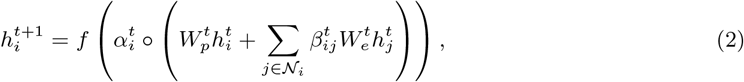

where *f* is a nonlinearity function and ⚬ is the element-wise product, 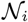 is the set of neighbors of node *i* (not including itself), 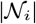 is the number of neighbors of node *i*. We use an exponential linear unit (ELU) for *f*. As in (Veličković et al., 2018), we use *K* heads and concatenate the features as

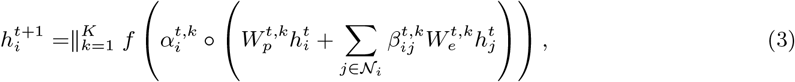

where 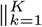 means concatenation of *K* independent heads. Therefore, 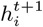 will have *KF′* features for *i* ∈ {1, …, *N*} and *t* ∈ {1, …, *T*}, where *T* is the number of GAT layers.

As we see in equation (3), the updated feature 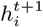 at bin *i* is also a function of the number of its neighbors, namely 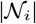. The impact of node cardinality on its feature is nontrivial by adding 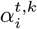 in (3). We mentioned that Epi-GraphReg models are cell-type agnostic, meaning that they can be generalized from one cell type to another one. When we want to train the Epi-GraphReg models on some cell types and test them on other unseen cell types, this might lead to performance degradation if the distributions of node cardinality of graphs in the train and test cells are different. This could happen for many reasons, including the library sizes or FDR levels of Hi-C and HiChIP data from which we extract the graphs. For example, in our experiments, we noticed this discrepancy between the graphs of GM12878 and K562 cells. In order to solve this and improve the generalization performance, we normalize the attention weights of each node *i* such that the summation of all the attention weights becomes one and we remove the cardinality parameter 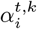. As such, the normalized updates of GAT layers can be written as

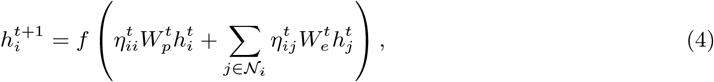

where 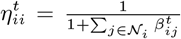 is the attention coefficient of the node *i* (or promoter coefficient) and 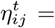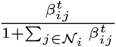 is the attention coefficient from the node *j* to the node *i* (or enhancer coefficient). We see that *η_ij_* ∈ [0, 1] for all *i, j* ∈ {1, …, *N*} and all the attention coefficients of each node *i* sum up to one: 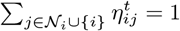. Similar to (3), when having K heads, the normalized updates would be

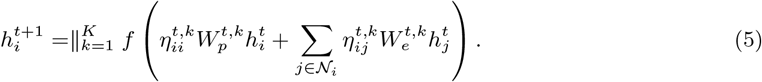

### Poisson regression and loss function

After the GAT layers in GraphReg models, the last layer before the predictions is a width-1 CNN layer (equivalent to fully connected layer) with exponential nonlinearity in order to predict the CAGE value at each bin. As the CAGE values are counts, we used Poisson regression, meaning that the expected value of each (CAGE) output given the inputs is the mean of a Poisson distribution.

For the Epi-GraphReg model, we let *X* denote the 1D epigenomic inputs for a 6Mb region, *G* the corresponding graph, *Y* = [*y*_1_, …, *y_N_*] the observed CAGE signal across 5Kb bins, and 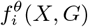 the predicted CAGE signal for the bin *i*, where *θ* represents parameters of the model. Now we assume that *y_i_*|*X, G* ~ Poisson(*λ_i_*), where 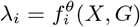 and *E*(*Y* |*X, G*) = *f^θ^*(*X, G*). Hence, the loss function is the negative log-likelihood of the Poisson distribution,

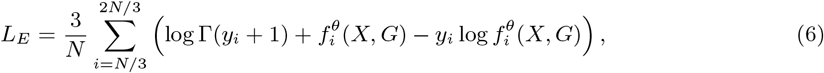

where Γ(.) is the gamma function. Note that we train our model for the middle one third region (2Mb or middle *N*/3 bins) instead of the whole 6Mb region (all *N* bins) in each batch. Since we include 3D interactions of genomic distance up to 2Mb, promoters in the middle 2Mb region can see the effects of distal enhancers in the full 6Mb input region. In the test phase we also restrict to predictions on the middle 2Mb regions. We shift input regions by 2Mb across genome, so there is no overlap between predictions in two different batches. Note that depending on the number of GAT layers used, we can see the effects of enhancers via multi-hop interactions up to 4Mb away from the promoters. If we have *L* GAT layers, message passing is done *L* times over the graph, and the promoters will see the effects of enhancers up to *L* hops.

For the Seq-GraphReg model, we also predict the 1D epigenomic data with a resolution of 100bp after the dilated CNN layers from DNA sequence. Here we let *X* denote the DNA sequence of the 6Mb input region, *G* the corresponding graph, 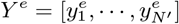 the observed 100bp epigenomic inputs (over *N*′ = 60000 bins), and 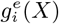 the predicted epigenomic signal *e* for the bin *i*. The loss for each epigenomic signal *e* is given by

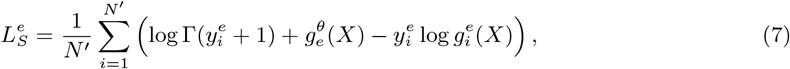

and the CAGE loss 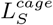, similar to (6), can be written as

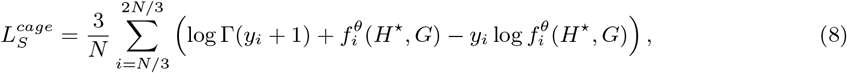

where *H*^⋆^ is the bottleneck representation just before dilated CNN layers (shown by star (⋆) in **Supplemental Fig. S1**) at 100bp resolution. The final loss function to be minimized is the convex combination of the CAGE loss and the epigenomic losses,

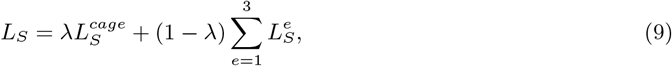

where *λ* is a hyperparameter to control the weights of the CAGE loss versus the epigenomic losses. We use a sigmoid-like schedule for *λ* given by 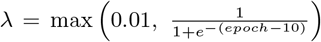, which starts with a small value *λ* = 0.01 and approaches a value close to one at higher epochs. This schedule lets the model first learn meaningful motif representations from DNA sequences in order to predict the epigenomic data and then puts higher weights on predicting the CAGE signals from those motif representations at 100bp resolution. *NLL* in all the figures of the paper refers to *L_E_* and 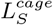 for epigenome-based and sequence-based models, respectively.

### End-to-end versus separate training of Seq-GraphReg

Seq-GraphReg as shown in **Supplemental Fig. S1** is an end to end model with two main tasks: (1) epigenomic tracks are predicted from DNA sequences; and (2) CAGE is predicted from the bottleneck representation *H*^⋆^ learned in first task. For end-to-end training, we need to load the entire 6Mb DNA sequence to the GPU memory, and this can limit our capability to use more CNN filters in the first layers to learn a richer bottleneck representation. Therefore, the overall CAGE prediction might be sacrificed because of the GPU memory limitations. In order to resolve this problem, we propose another scheme for training Seq-GraphReg, to which we refer as separate training. The separate training works as follows. First, we perform task (1) to predict the epigenomic data from DNA sequences of length 100Kb. Then, we freeze the learned parameters of the CNN layers and get the bottleneck representations *H*^⋆^, which are at 100bp resolution. Finally, we use *H*^⋆^ as the input to the second block (**Supplemental Fig. S1**) to predict the CAGE values. The second block has several CNN and maxpool layers to bring the resolution to 5Kb, and then it is given to 3 GAT layers to predict the CAGE values.

We have shown that separate training on K562 outperforms end-to-end training in terms of CAGE prediction performance. The reason for the improvement is that in separate training, we use 100Kb DNA sequences (instead of 6Mb), so we do not exhaust GPU memory and so can use more CNN filters to learn richer bottleneck representations which consequently lead to superior gene expression prediction. However, this prediction improvement comes at the cost of losing base-pair level saliency scores via backpropagation. The reason for this is that in order to get the gradient of the output CAGE with respect to the input DNA sequence via backpropagation, we need an end-to-end model from input to the output. Nevertheless, we can use perturbation-based feature attribution methods such as in silico mutagenesis (ISM) for the separately trained Seq-GraphReg models (as we have done in **Fig. 4E**). However, backpropagation-based feature attribution methods, such as gradient-by-input and DeepSHAP, are much faster than ISM. Overall, there is a trade-off between better prediction performance (achieved by separate training) and having fast backpropagation-based feature attribution capability (achieved by end-to-end training).

### Dilated CNN layers and FFT loss in Seq-GraphReg

In separate training of Seq-GraphReg, we also studied the effects of using dilated CNN layers and also adding FFT loss to get better motif interpretations. In task (1) of separate training, where we predict epigenomic data from DNA sequence, we removed the residual dilated CNN layers and instead we used three CNN layers after the bottleneck *H*^⋆^ for each task of predicting H3K4me3, H3K27ac, and DNase. We observed that not using dilated layers in fact helped prediction performance in K562 (**Supplemental Fig. S8**). This is partly because when we do not use dilated layers, the CNN filters can focus more on local peak regions and learn better motif representations.

As our goal is not only prediction but also interpretation of base-pair feature attribution methods, we used the idea of adding FFT loss to the prediction loss (NLL) at training, as suggested in (Tseng et al., 2020). Here we explain how we used FFT loss in separate training of Seq-GraphReg models. In task (1), let *g*(*H*^⋆^, *X*) be the gradient of bottleneck *H*^⋆^ with respect to the input DNA sequence *X* and summed over 4 bases as our attribution vector. So, the length of the vector *g*(*H*^⋆^, *X*) is the same as that of *X*. Let *a* denote the magnitudes of the positive-frequency Fourier components of (a slightly smoothed version of) *g*(*H*^⋆^, *X*), and 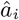 denote the *i*-th component of 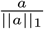 (smaller i correspond to lower frequencies). Similar to (Tseng et al., 2020), we penalize the high-frequency components as:

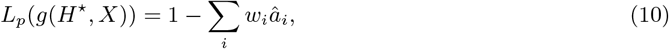

where *w_i_* = 1 for *i* ≤ *T*, and 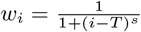 for *i > T*. The total loss to be trained is 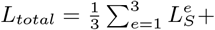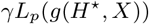, where 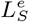 is the prediction loss (NLL) of the epigenomic data *e*, *L_p_*(*g*(*H*^⋆^, *X*)) is attribution prior, and *γ* is a positive scalar to control the contribution of attribution loss. We have used *s* = 0.2 and *γ* = 1 as suggested in (Tseng et al., 2020). We have applied ISM to the Seq-GraphReg models trained separately without dilation and with FFT loss and observed that relevant TF motifs can be identified in distal enhancer regions (**Fig. 4E**).

### Model architecture and training

For Epi-GraphReg and Seq-GraphReg we used respectively 2 and 3 GAT layers with 4 heads in each layer (**Supplemental Fig. S1**). For Epi-CNN and Seq-CNN we used 8 dilated CNN layers to increase their receptive field up to 2.5Mb. For human cell lines GM12878, K562, and hESC, we held out chromosomes *i* and *i* + 10 for validation and chromosomes *i* + 1 and *i* + 11 for test, for *i* = 1, …, 10, and trained on all remaining chromosomes except X and Y. For mouse cell line mESC, we held out chromosomes *i* and (*i* + 10 mod 20) + sign(*i* ≥ 10) for validation and chromosomes *i* + 1 and (*i* + 11 mod 20) + sign(*i* ≥ 9) for test, for *i* = 1, …, 10, and trained on all remaining chromosomes except X and Y.

### TF knockout analysis and motif ablation experiments

For RNA-seq data sets from CRISPRi TF KO experiments, we performed differential gene expression analyses using DESeq2 (Anders and Huber, 2010) between control and TF KO K562 cells for each TF. We only considered TFs whose KO led to significant downregulation (adjusted *p* < 0.05) of at least 200 genes in order to have robust statistics. To assess the ability of Seq-GraphReg and Seq-CNN to predict the downregulation of genes as direct effects of TF KO, we performed an in silico KO of each TF by zeroing out all the motifs for the TF in 6Mb DNA sequences encompassing candidate target genes. Then we used the models to predict the change in gene expression caused as direct effects of each TF KO. Similar to the true KO experiments, we used two trained models each for Seq-GraphReg and Seq-CNN as two replicates to predict control and in silico KO expression levels. We then performed differential expression analysis using DESeq2 to get the predicted logFC (log fold change) and adjusted *p*-values. We ranked the significantly regulated genes (adjusted *p* < 0.05) with true CAGE values at least 20 based on predicted logFC, and used the 100 genes with largest negative predicted logFC as the predicted target genes for each model.

## Code availability

The code to preprocess data, train models, and perform the analyses in the paper, as well as all the trained models are available in the supplemental material and on https://github.com/karbalayghareh/GraphReg.

## Data

We downloaded the epigenomic and CAGE data of GM12878, K562, hESC, and mESC cells from ENCODE (https://www.encodeproject.org/) portal with the following accession numbers: K562 CAGE: ENCFF623BZZ and ENCFF902UHF; K562 DNase-seq: ENCFF826DJP; K562 H3K4me3: ENCFF689TMV; K562 H3K27ac: ENCFF010PHG; K562 JUND TF ChIP-seq: ENCFF709JGL; GM12878 CAGE: ENCFF915EIJ and ENCFF990KLZ; GM12878 DNase-seq: ENCFF775ZJX and ENCFF783ZLL; GM12878 H3K4me3: ENCFF818GNV; GM12878 H3K27ac: ENCFF180LKW; hESC (H1) CAGE: ENCFF525HJR; hESC (H1) DNase-seq: ENCFF131HMO; hESC (H1) H3K4me3: ENCFF760NUN; hESC (H1) H3K27ac: ENCFF919FBG; mESC (ES-E14) DNase-seq: ENCFF754ILF and ENCFF785XJZ; mESC (ES-E14) H3K4me3: ENCFF240MDV; mESC (ES-E14) H3K27ac: ENCFF163HEV. We downloaded mESC CAGE data from GEO with the accession numbers GSM3852792, GSM3852793, and GSM3852794. We got the H3K27ac HiChIP data of K562, GM12878, and mESC from GEO with the the accession numbers GSE101498, GSE101498, and GSE113339, respectively. We obtained Hi-C data of K562 and GM12878 from GEO with the accession number GSE63525. We obtained Micro-C data of hESC (H1) from 4DN portal with the accession number of 4DNFI2TK7L2F. We got the CRISPRi FlowFISH enhancer validation data set for K562 from Supplemental Table 6a of (Fulco et al., 2019). We got the CRISPRi TAP-seq enhancer validation data set for K562 (chromosomes 8 and 11) from Supplemental Tables 2 and 3 of (Schraivogel et al., 2020). We acquired ABC enhancer predictions of the K562 (at chromosomes 8 and 11) from https://osf.io/f2uvz/. We downloaded CRISPRi RNA-seq (transcription factor (TF) knockout) experiments of 51 TFs (plus 2 controls) in K562 from ENCODE with the accession numbers ENCSR439NIR, ENCSR447WYJ, ENCSR490XKT, ENCSR895NMN, ENCSR747UPR, ENCSR045BVL, ENCSR658NCC, ENCSR685TOO, ENCSR157JAK, ENCSR290RLW, ENCSR849RWU, ENCSR177JPA, ENCSR434ZAM, ENCSR467RQJ, ENCSR179XMY, ENCSR231XYT, ENCSR569WHC, ENCSR612JVF, ENCSR844UQZ, ENCSR466OEJ, ENCSR156EPV, ENCSR991TCB, ENCSR559VHS, ENCSR109KMO, ENCSR539CHL, ENCSR153WDD, ENCSR046XJC, ENCSR692NVT, ENCSR932JIP, ENCSR315EHR, ENCSR612NHI, ENCSR143YTV, ENCSR997FLI, ENCSR588MYR, ENCSR366ONT, ENCSR708MPN, ENCSR747ZDR, ENCSR521VAG, ENCSR045LNG, ENCSR199FFQ, ENCSR208KNW, ENCSR004ZBD, ENCSR975MRK, ENCSR966USM, ENCSR496MYK, ENCSR866PAO, ENCSR205DTM, ENCSR212QDE, ENCSR349LHQ, ENCSR043UEE, ENCSR785ATE, ENCSR095PIC, ENCSR016WFQ.

## Acknowledgement

This work was supported by NIH/NHGRI U01 award HG009395 and NIH/NIDDK U01 award DK128852.

## Author Contributions

AK developed the machine learning models, performed all computational experiments, and co-wrote the manuscript. MS performed analysis on HiChIP data sets. CL supervised the research and co-wrote the manuscript.

