## Supplemental Material for "Chromatin interaction aware gene regulatory modeling with graph attention networks"

| | Uses DNA sequence input? | Uses epigenomic input? | Uses 3D conformation input? | Uses CNNs? | Uses GNN? | Receptive field from TSS | Captures promoter-proximal enhancers? | Captures distal (> 100Kb) enhancers? | Captures promoter-proximal TFs? | Captures distal (> 100Kb) TFs? |
|---|---|---|---|---|---|---|---|---|---|---|
| Enformer | ✓ | ✗ | ✗ | ✓ | ✗ | 100 Kb | ✓ | ✗ | ✓ | ✗ |
| Basenji | ✓ | ✗ | ✗ | ✓ | ✗ | 32 Kb | ✓ | ✗ | ✓ | ✗ |
| Expecto | ✓ | ✗ | ✗ | ✓ | ✗ | 20 Kb | ✓ | ✗ | ✓ | ✗ |
| Xpresso | ✓ | ✗ | ✗ | ✓ | ✗ | 7 Kb up<br>3.5 Kb down | ✓ | ✗ | ✓ | ✗ |
| GC-MERGE | ✗ | ✓ | ✓ | ✗ | ✓ | NA | ✗ | ✗ | ✗ | ✗ |
| DeepExpression | ✓ | ✗ | ✓ | ✓ | ✗ | 1 Mb | ✗ | ✗ | ✓ | ✗ |
| **GraphReg** | ✓ | ✓ | ✓ | ✓ | ✓ | 2-4 Mb | ✓ | ✓ | ✓ | ✓ |

Table S1: Deep learning methods for gene expression prediction.

| | K562 | GM12878 |
|---|---|---|
| Epi-GraphReg | **0.885** | **0.884** |
| Epi-CNN | 0.879 | 0.877 |
| GC-MERGE* | 0.76 | 0.73 |

Table S2: Pearson correlation (R) of true versus predicted gene expression in epigenome-based models. ∗The results of GC-MERGE are reported from their paper.

| | K562 | mESC | GM12878 |
|---|---|---|---|
| Seq-GraphReg | **0.743** | **0.835** | **0.721** |
| Seq-CNN | 0.655 | 0.829 | 0.580 |
| DeepExpression* | 0.68 | 0.81 | - |
| Xpresso* | 0.714 | 0.768 | - |
| Basenji | 0.566 | - | 0.697 |

Table S3: Pearson correlation (R) of true versus predicted gene expression in sequence-based models. ∗The results of DeepExpression and Xpresso are reported from their papers. We ran Basenji on K562 and GM12878 individually with four tracks (CAGE, DNase, H3K27ac, and H3K4me3) ten times with the same valid/test/train chromosome splits.

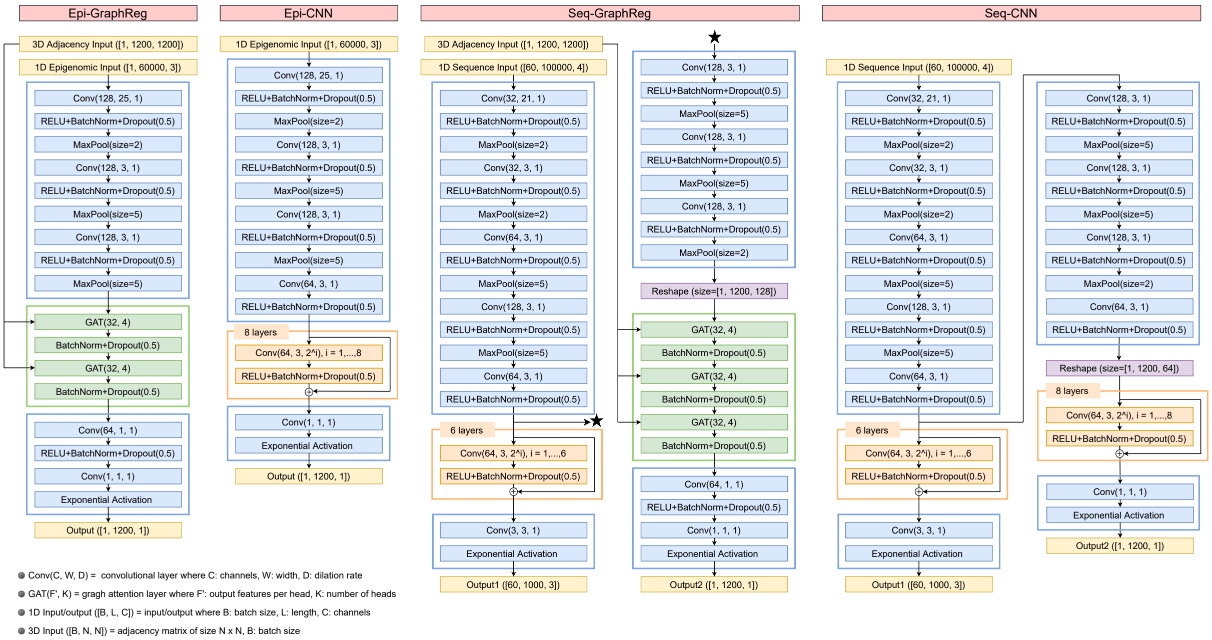


Figure S1: **Architectures of Epi-GraphReg, Epi-CNN, Seq-GraphReg, and Seq-CNN models**. Blue blocks show the CNN layers: Conv(C, W, D) is a CNN layer with C channels, width W, and dilation rate D. Green blocks show the GAT layers: GAT(F′, K) is a GAT layer with F′ output features per head and K heads; hence, GAT(F′, K) layer will have F′K output features overall. Orange blocks show the residual dilated CNN layers, where the input and output of each layer are summed. For the CNN models, for a fair comparison with GraphReg models, we have increased the receptive fields up to 2.5Mb upstream and downstrem of TSS bins by using 8 dilated layers whose dilation rate is multiplied by 2 at each layer. Yellow blocks show the inputs and outputs of the models. Output([B, L, C]) and input([B, L, C]) show the predicted outputs of the models and their 1D inputs, where B denotes the batch size, L is the length, and C is the number of channels. 3D input([B, N, N]) denotes B (batch size) adjacency matrix of size N×N derived from H3K27ac HiChIP, Hi-C, or Micro-C data at the resolution of 5Kb for a 6Mb genomic region (hence, N=1200). In the Seq-GraphReg and Seq-CNN models, there are two types of outputs: output1 denotes the predicted epigenomic data (H3K4me3, H3K27ac, DNase) at 100bp resolution and output2 denotes the predicted CAGE at 5Kb resolution. In the separate training of Seq-GraphReg, as opposed to end-to-end training, output1 is first predicted from DNA sequence, then the bottleneck representation (shown by star) is given to the GAT block to predict CAGE.

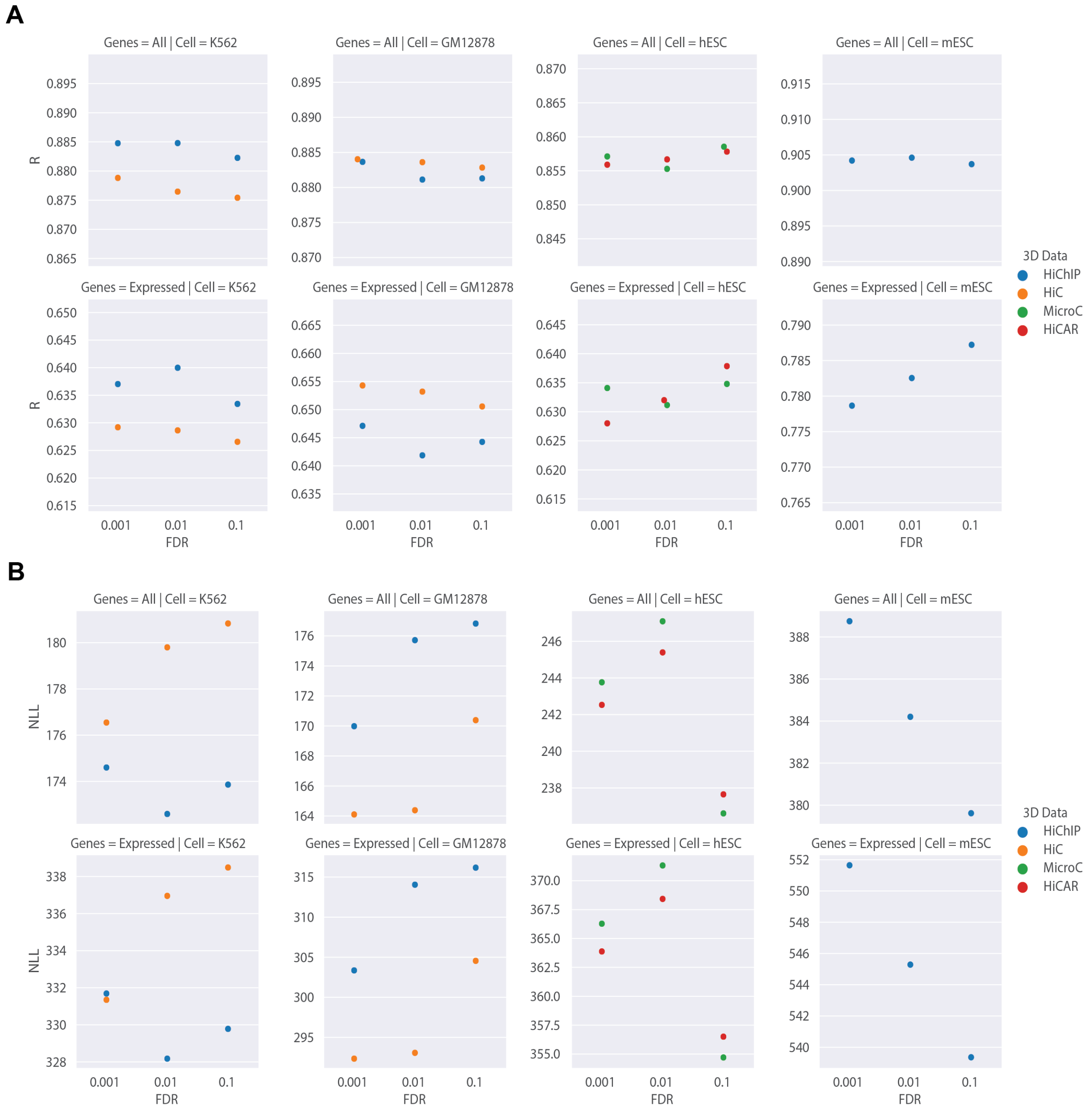


Figure S2: **The effects of 3D data and FDR thresholds on Epi-GraphReg. A.** The Pearson correlation (R) of the log-normalized ($\log(x+1)$) true and predicted gene expression values for the cell lines K562, GM12878, hESC, and mESC, using graphs extracted from different 3D data such as Hi-C, HiChIP, Micro-C, and HiCAR, with three different levels of FDR at 0.1, 0.01, and 0.001. **B.** The same experiments reporting NLL.

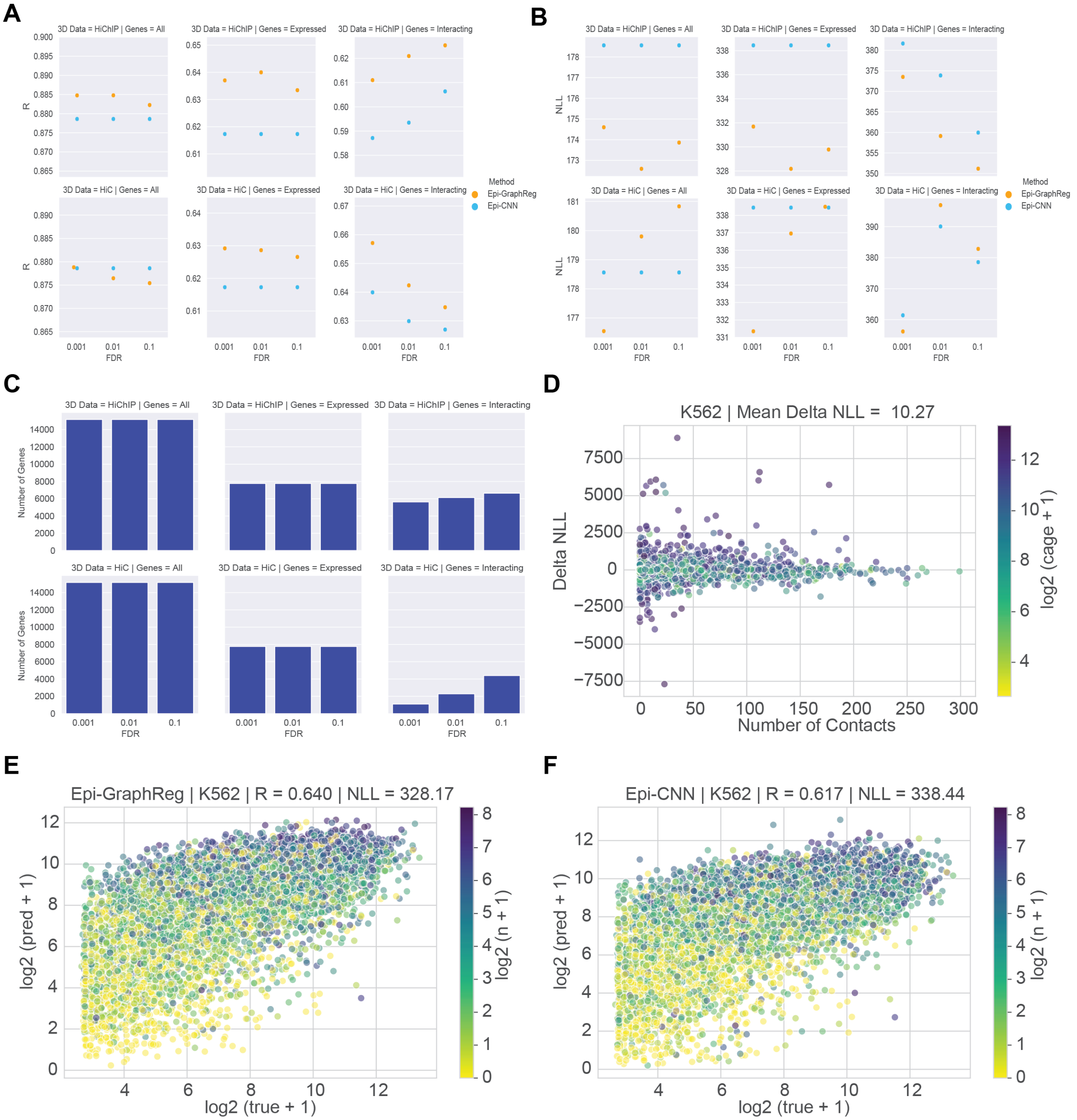


Figure S3: **Epi-GraphReg predictions in K562. A.** The Pearson correlation (R). **B.** Negative log-likelihood (NLL). **C.** Number of the test genes (concatenated from 20 test chromosomes from 10 runs of the models each having 2 test chromosomes). **D.** Difference in NLL (Delta NLL) of Epi-GraphReg (HiChIP with FDR 0.01) and Epi-CNN versus number of contacts for each gene. If Delta NLL is positive, the prediction of Epi-GraphReg for that gene is better than Epi-CNN. **E.** Scatter plot (true versus predicted CAGE) of Epi-GraphReg (HiChIP with FDR 0.01). **F.** Scatter plot (true versus predicted CAGE) of Epi-CNN. All scatter plots are for expressed genes.

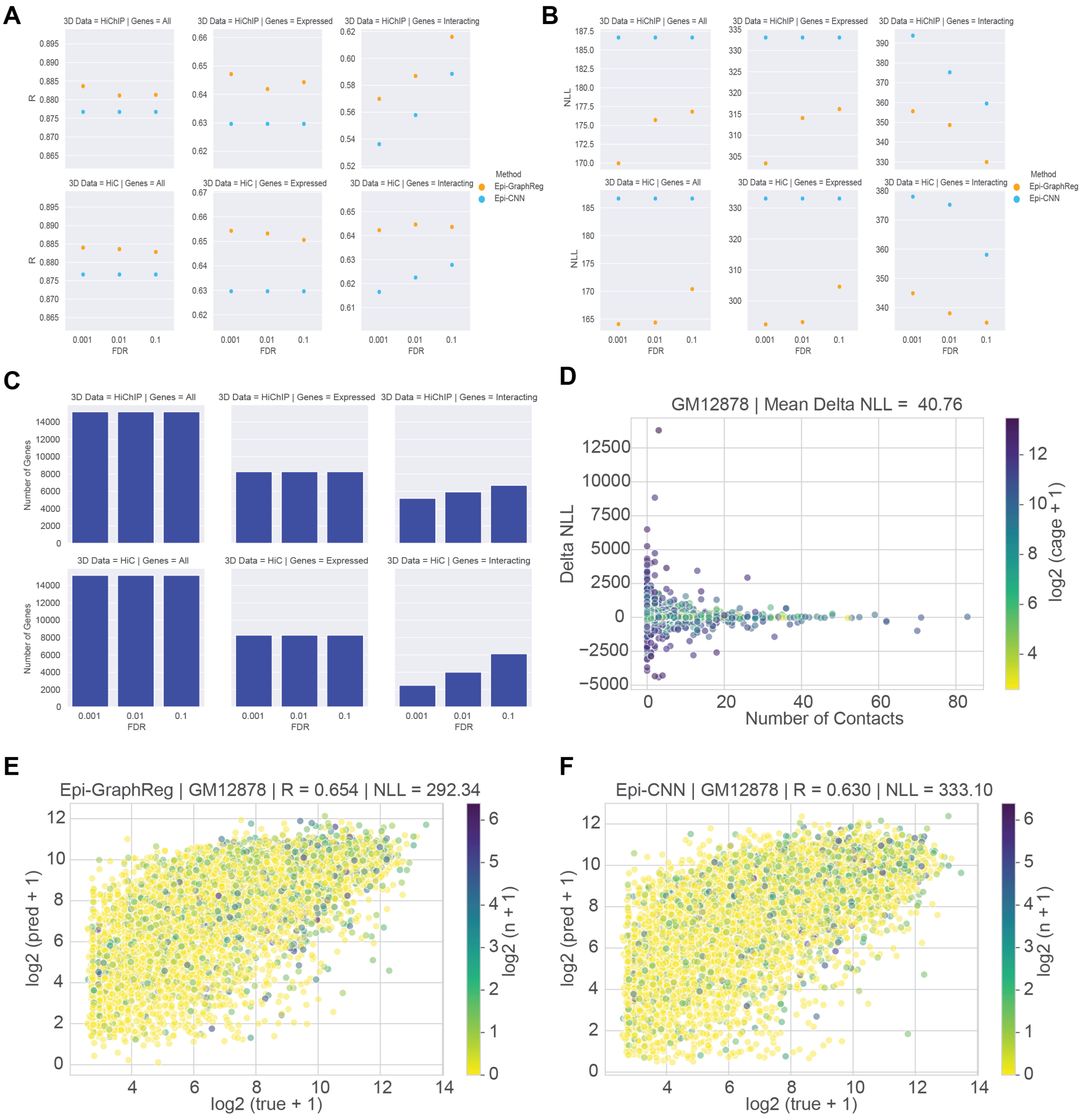


Figure S4: **Epi-GraphReg predictions in GM12878. A.** The Pearson correlation (R). **B.** Negative log-likelihood (NLL). **C.** Number of the test genes (concatenated from 20 test chromosomes from 10 runs of the models each having 2 test chromosomes). **D.** Difference in NLL (Delta NLL) of Epi-GraphReg (Hi-C with FDR 0.001) and Epi-CNN versus number of contacts for each gene. If Delta NLL is positive, the prediction of Epi-GraphReg for that gene is better than Epi-CNN. **E.** Scatter plot (true versus predicted CAGE) of Epi-GraphReg (Hi-C with FDR 0.001). **F.** Scatter plot (true versus predicted CAGE) of Epi-CNN. All scatter plots are for expressed genes.

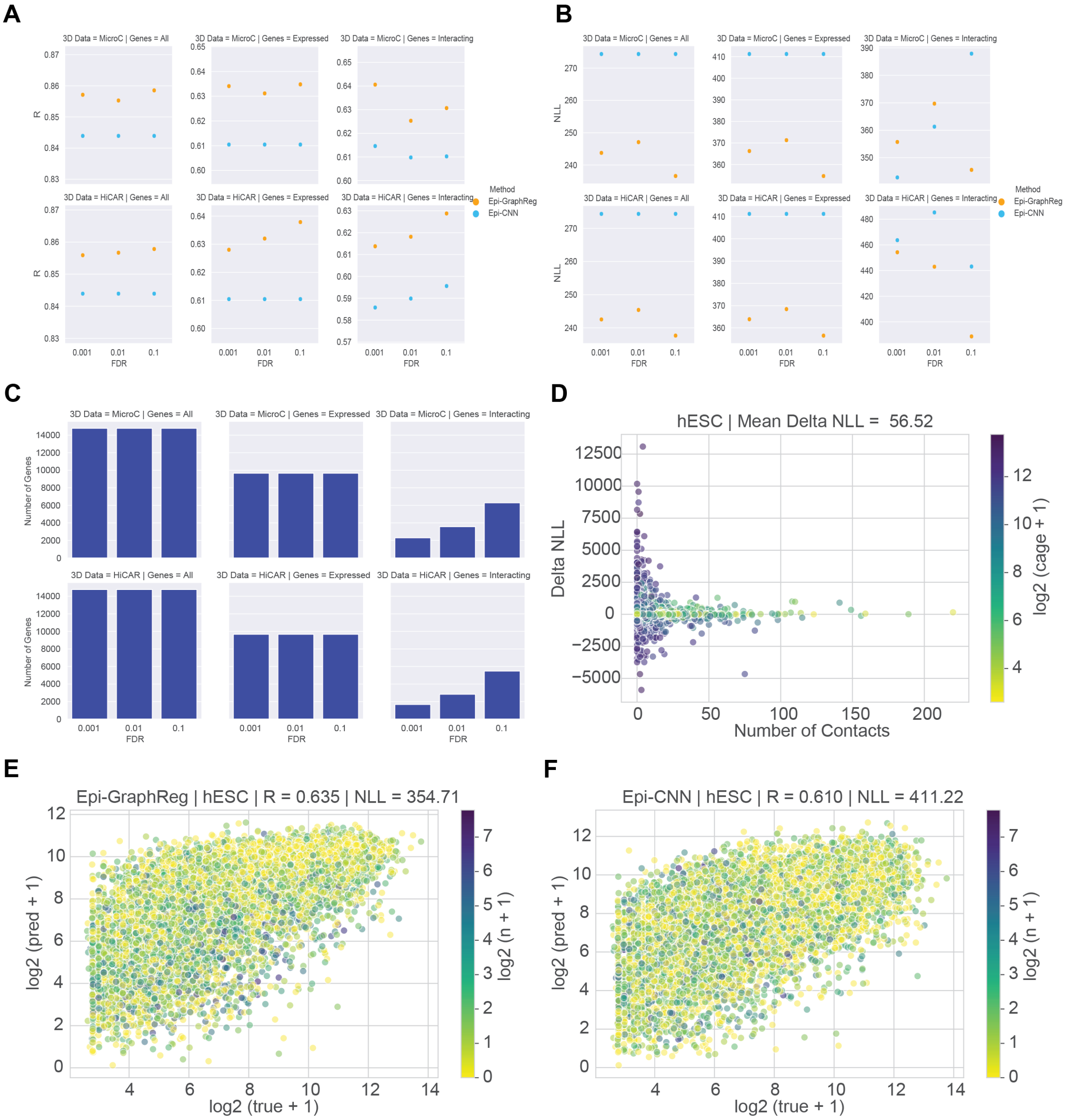


Figure S5: **Epi-GraphReg predictions in hESC. A.** The Pearson correlation (R). **B.** Negative log-likelihood (NLL). **C.** Number of the test genes (concatenated from 20 test chromosomes from 10 runs of the models each having 2 test chromosomes). **D.** Difference in NLL (Delta NLL) of Epi-GraphReg (Micro-C with FDR 0.1) and Epi-CNN versus number of contacts for each gene. If Delta NLL is positive, the prediction of Epi-GraphReg for that gene is better than Epi-CNN. **E.** Scatter plot (true versus predicted CAGE) of Epi-GraphReg (Micro-C with FDR 0.1). **F.** Scatter plot (true versus predicted CAGE) of Epi-CNN. All scatter plots are for expressed genes.

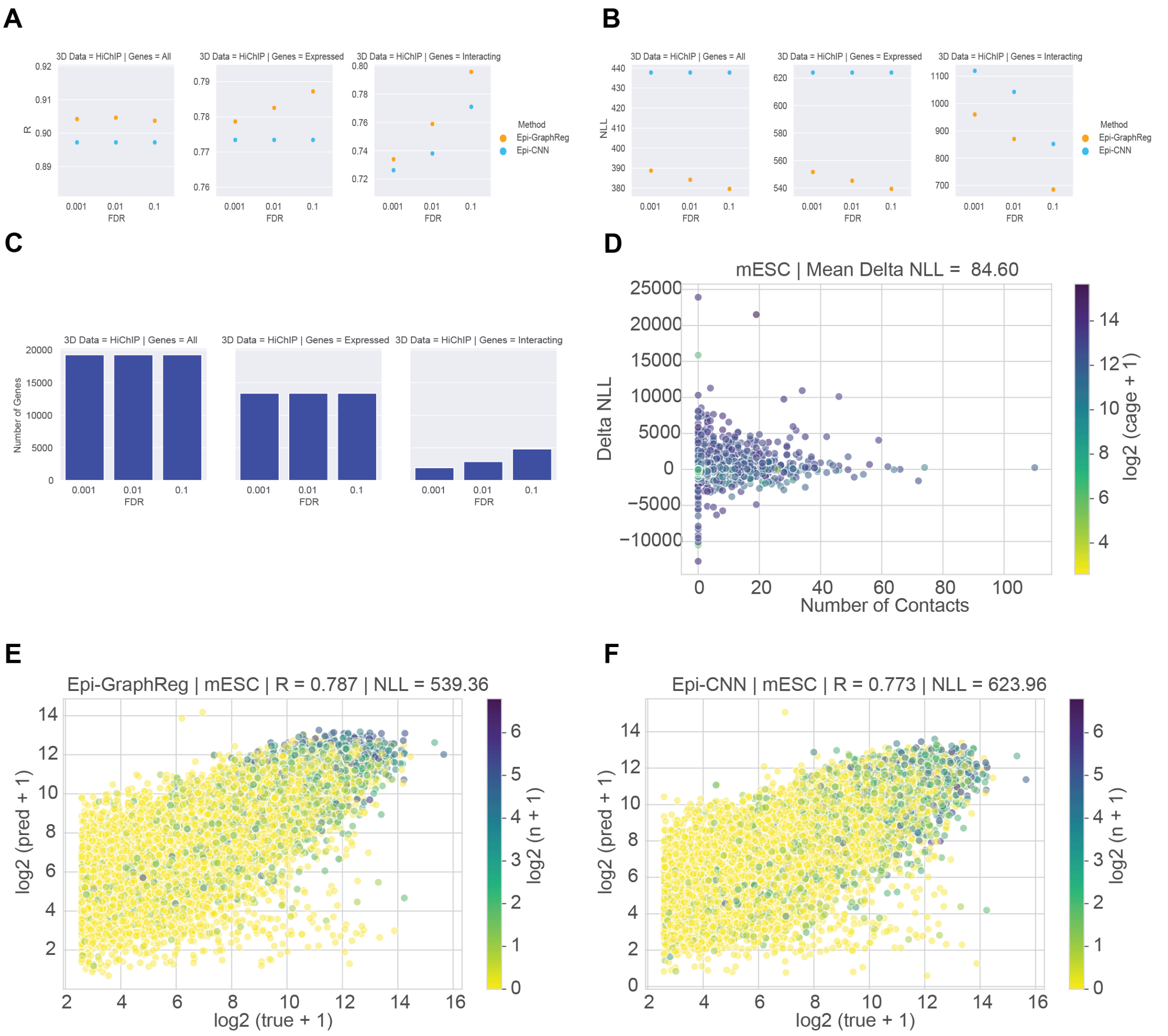


Figure S6: **Epi-GraphReg predictions in mESC. A.** The Pearson correlation (R). **B.** Negative log-likelihood (NLL). **C.** Number of the test genes (concatenated from 19 test chromosomes from 10 runs of the models each having 2 test chromosomes). **D.** Difference in NLL (Delta NLL) of Epi-GraphReg (HiChIP with FDR 0.1) and Epi-CNN versus number of contacts for each gene. If Delta NLL is positive, the prediction of Epi-GraphReg for that gene is better than Epi-CNN. **E.** Scatter plot (true versus predicted CAGE) of Epi-GraphReg (HiChIP with FDR 0.1). **F.** Scatter plot (true versus predicted CAGE) of Epi-CNN. All scatter plots are for expressed genes.

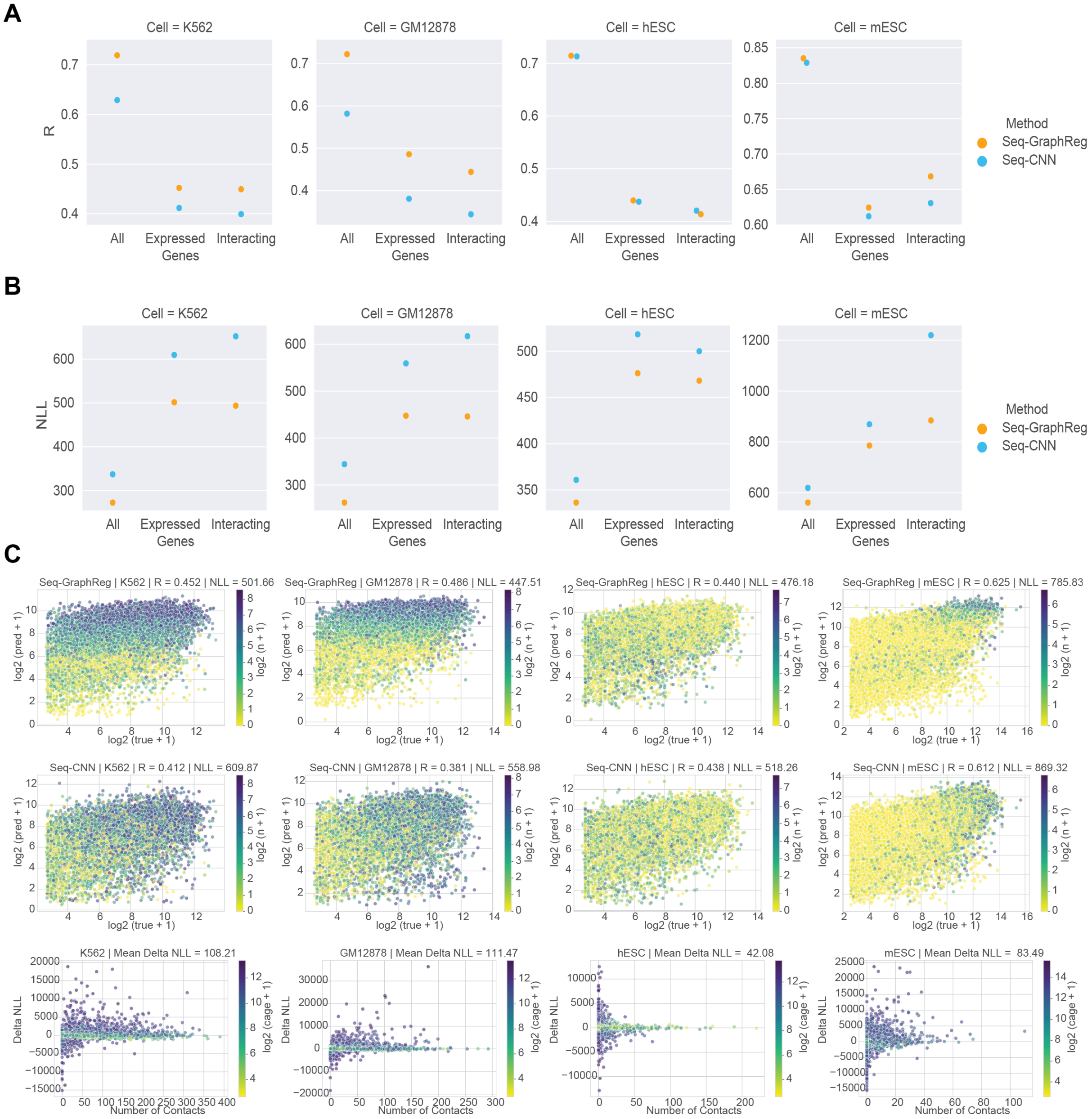


Figure S7: **End-to-end training of Seq-GraphReg and Seq-CNN in four different cell types.** HiChIP is used for K562, GM12878, and mESC, and Micro-C is used for hESC. We used FDR of 0.1 in all cell types. **A.** The Pearson correlation (R). **B.** Negative log-likelihood (NLL). **C.** Scatter plots for predictions of Seq-GraphReg, Seq-CNN, and Delta NLL in all four cell types. Each column shows one cell type.

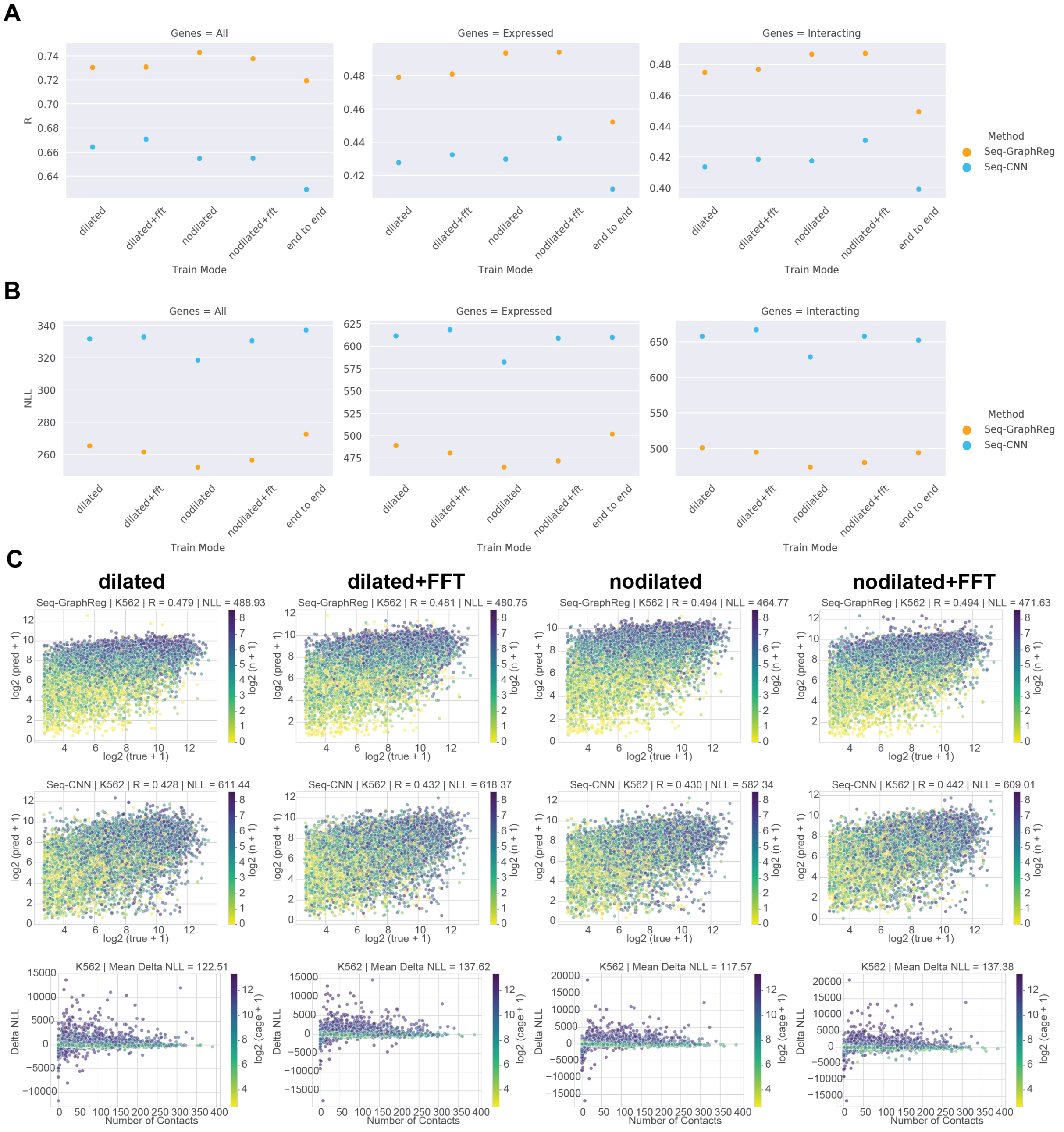


Figure S8: **Seq-GraphReg and Seq-CNN results in K562 with end-to-end and separate training.** Separate training means that instead of multitask learning of both epigenomic and CAGE-seq data (as in end-to-end training), first epigenomic data are predicted from DNA sequence, and then the bottleneck representation is used to predict CAGE. HiChIP data with FDR of 0.1 is used for all the experiments. The separate training has 4 versions: dilated, dilated+fft, nodilated, and nodilated+fft. **A.** The Pearson correlation (R). **B.** Negative log-likelihood (NLL). **C.** Scatter plots for predictions of Seq-GraphReg, Seq-CNN, and Delta NLL in each version of separate training.

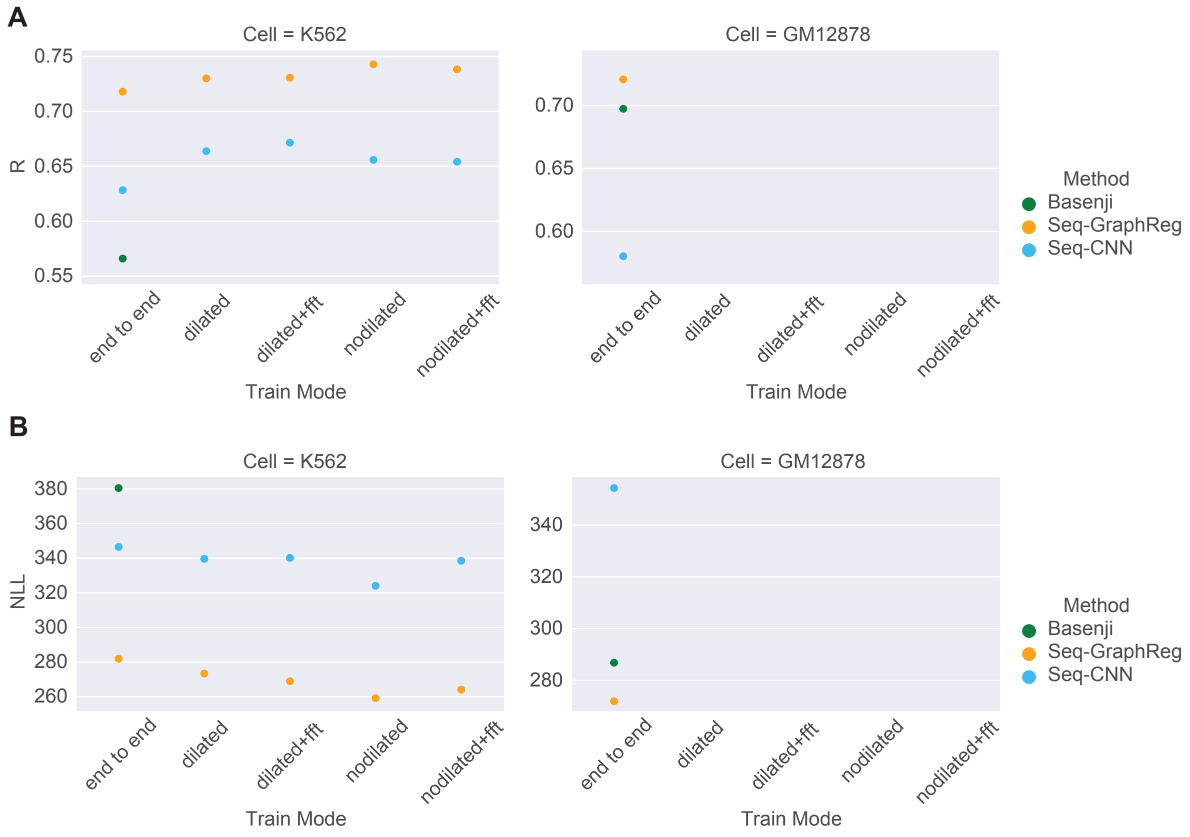


Figure S9: **Prediction performance of Seq-GraphReg, Seq-CNN, and Basenji on all genes concatenated from test chromosomes in cell types K562 and GM12878.** Seq-GraphReg and Seq-CNN are trained end-to-end and separately in K562 but only end-to-end in GM12878. Separate training in K562 includes four schemes: dilated, dilated with FFT, not dilated, and not dilated with FFT. HiChIP (FDR=0.1) is used for Seq-GraphReg. **A.** Pearson correlation (R), **B.** Negative log-likelihood (NLL).

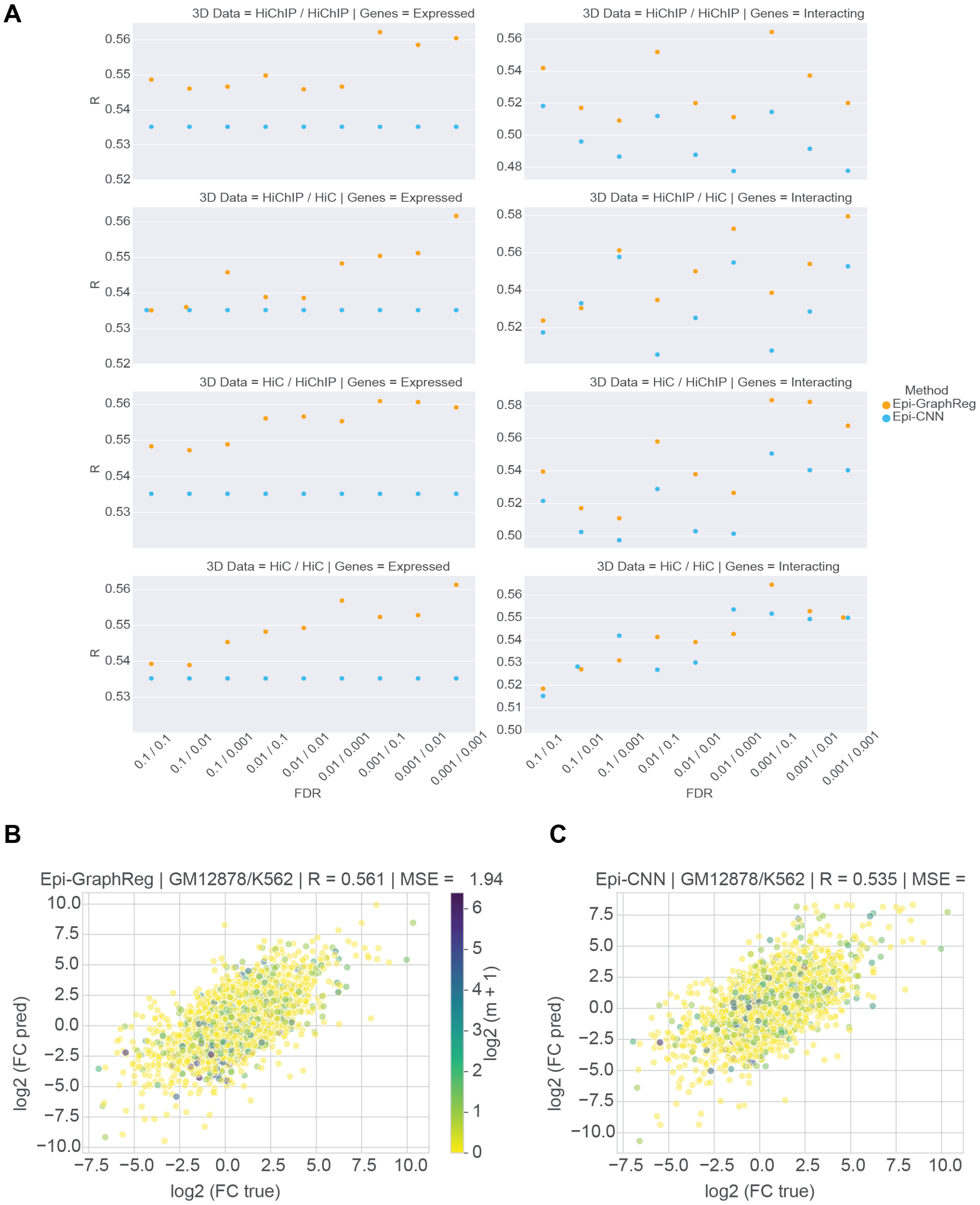


Figure S10: **Log fold change (logFC) prediction performance of Epi-GraphReg and Epi-CNN between GM12878 and K562. A.** Pearson correlation (R) between predicted and true logFC (GM12878/K562) in expressed and interacting genes for all combinations of 3D data and FDR values used in each cell type. Expressed and interacting genes here are the ones that are expressed (CAGE $\geq$ 5) and interacting in both GM12878 and K562. **B,C.** Scatter plots of predicted and true logFC for expressed genes (in both GM12878 and K562) by Epi-GraphReg and Epi-CNN, respectively, when using Hi-C (FDR=0.001) for GM12878 and HiChIP (FDR=0.01) for K562. $m$ shows the minimum number of 3D interactions in GM12878 and K562 in each TSS bin.

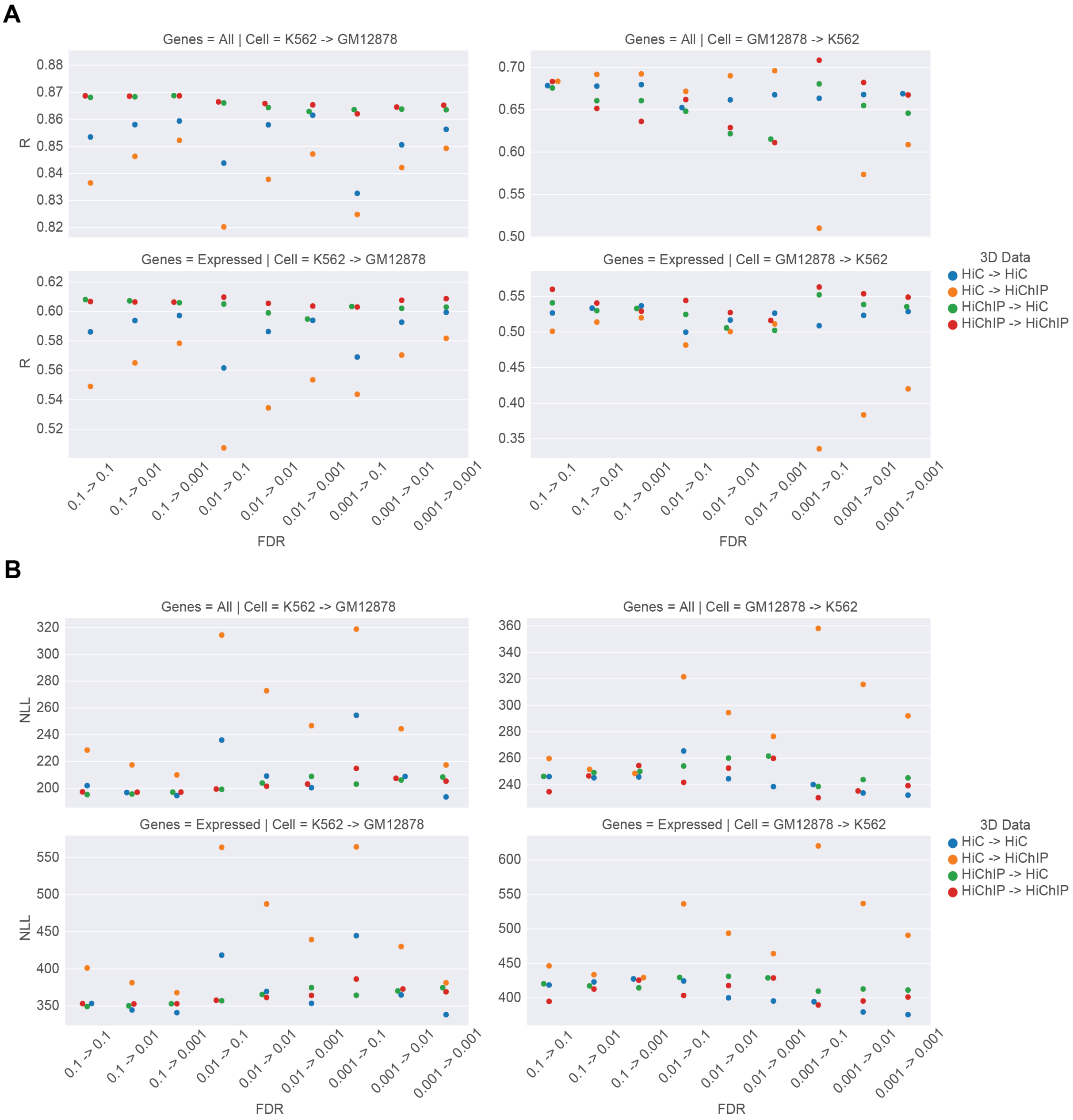


Figure S11: **Cross-cell-type generalization performance of Epi-GraphReg.** K562 and GM12878 cell types are used separately for training and test. These results are for cross-cell-type and cross-chromosome, meaning that the same test chromosomes in the first cell type (train cell type) are used as test chromosomes in the second cell type (test cell type). Hi-C and HiChIP data each with three FDR thresholds are used in both train and test cell types, so overall 36 combinations for each of K562 $\rightarrow$ GM12878 and GM12878 $\rightarrow$ K562. **A.** R in all and expressed genes of the test cell type. **B.** NLL in all and expressed genes of the test cell type.

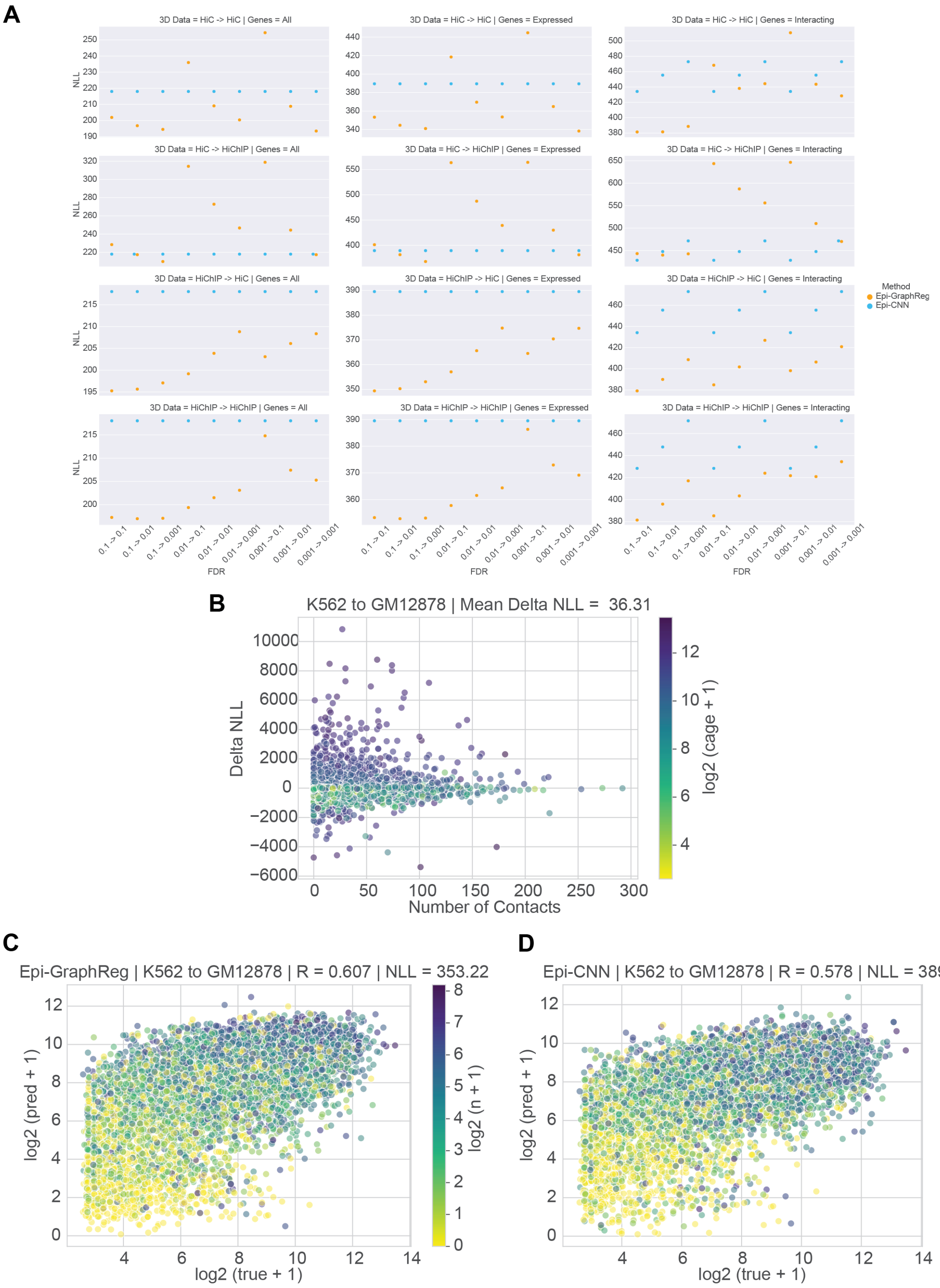


Figure S12: **Generalization performance from K562 to GM12878. A.** Negative log-likelihood (NLL) of Epi-GraphReg and Epi-CNN for all combinations of 3D data and FDRs in all, expressed, and interacting genes. **B, C, D.** Scatter plots for Delta NLL and predictions of Epi-GraphReg and Epi-CNN when using HiChIP (FDR=0.1) for both K562 and GM12878.

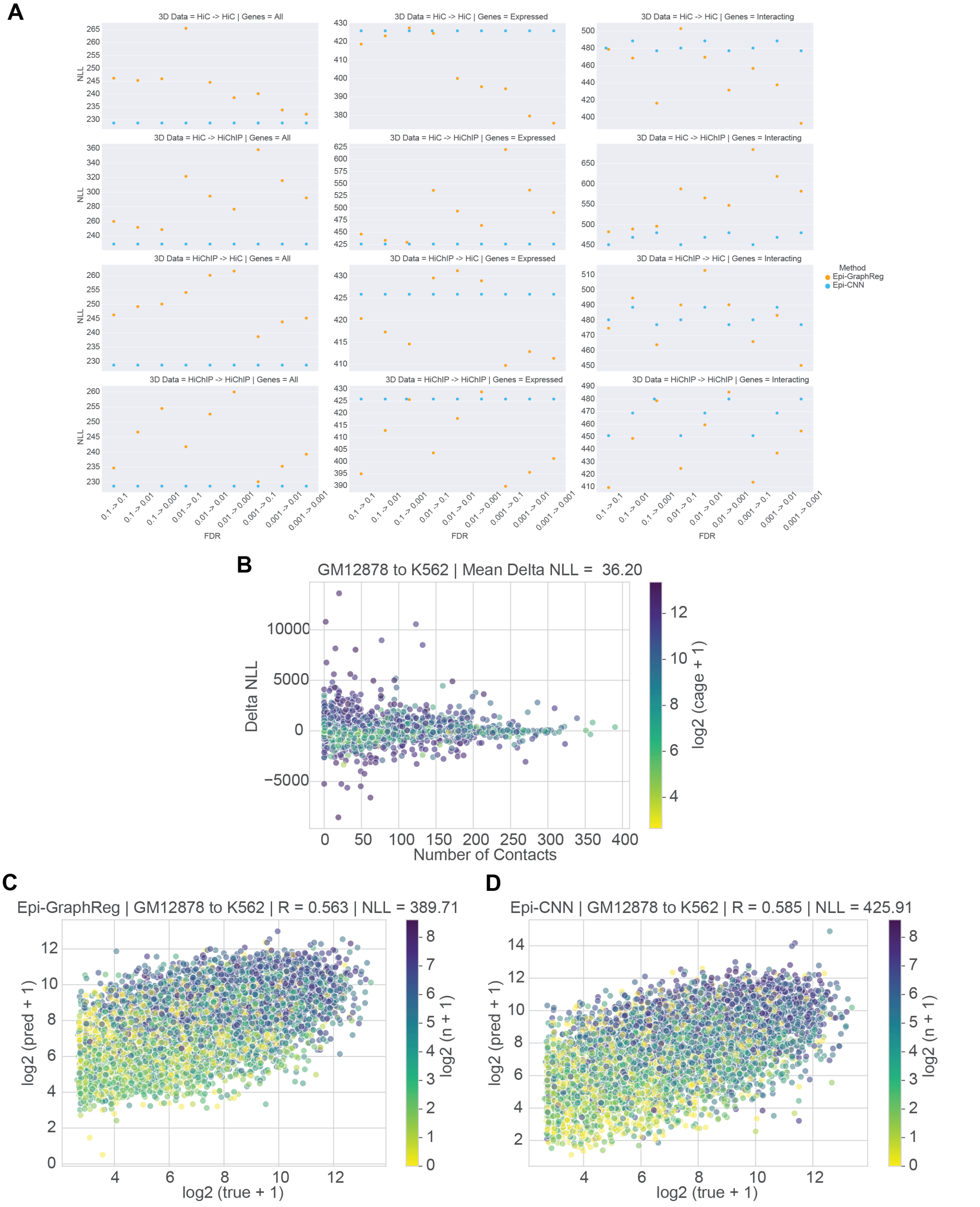


Figure S13: **Generalization performance from GM12878 to K562. A.** Negative log-likelihood (NLL) of Epi-GraphReg and Epi-CNN for all combinations of 3D data and FDRs in all, expressed, and interacting genes. **B, C, D.** Scatter plots for Delta NLL and predictions of Epi-GraphReg and Epi-CNN when using HiChIP (FDR=0.001) for GM12878 and HiChIP (FDR=0.1) for K562.

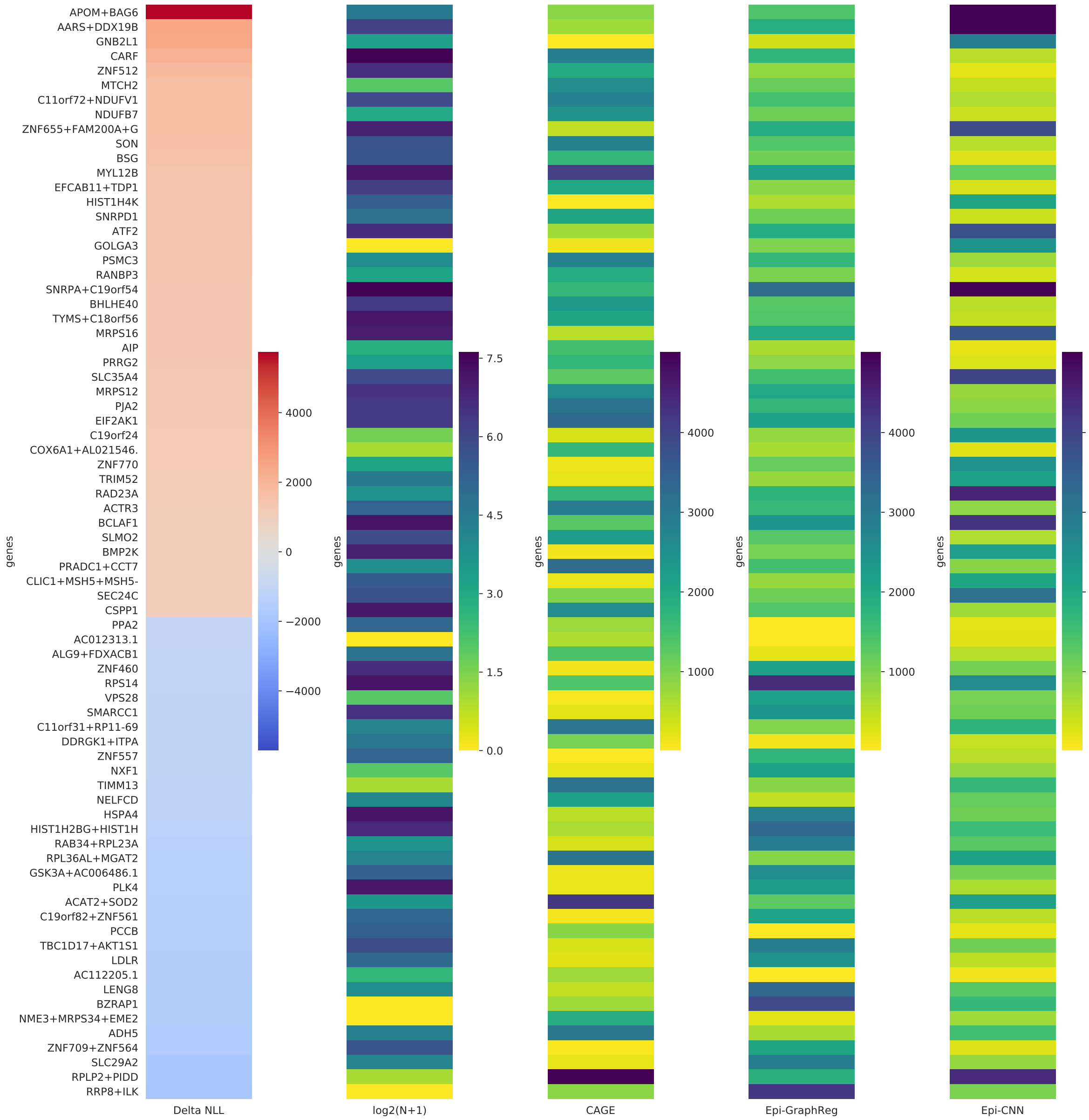


Figure S14: **K562 genes predicted accurately by either Epi-GraphReg or Epi-CNN and with significant difference in performance (Delta NLL)**. The results are for HiChIP (FDR=0.01).

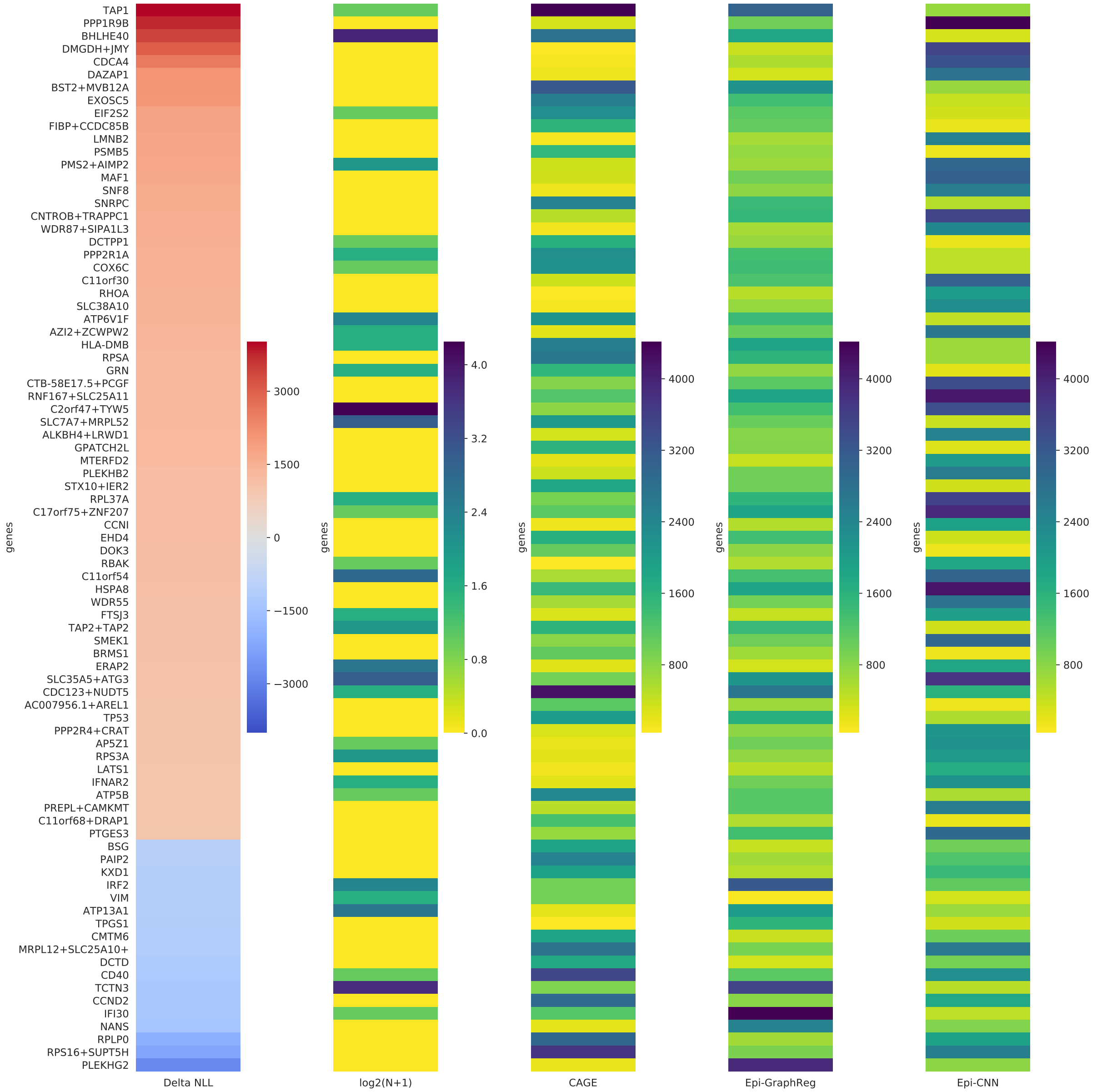


Figure S15: **GM12878 genes predicted accurately by either Epi-GraphReg or Epi-CNN and with significant difference in performance (Delta NLL)**. The results are for Hi-C (FDR=0.001).

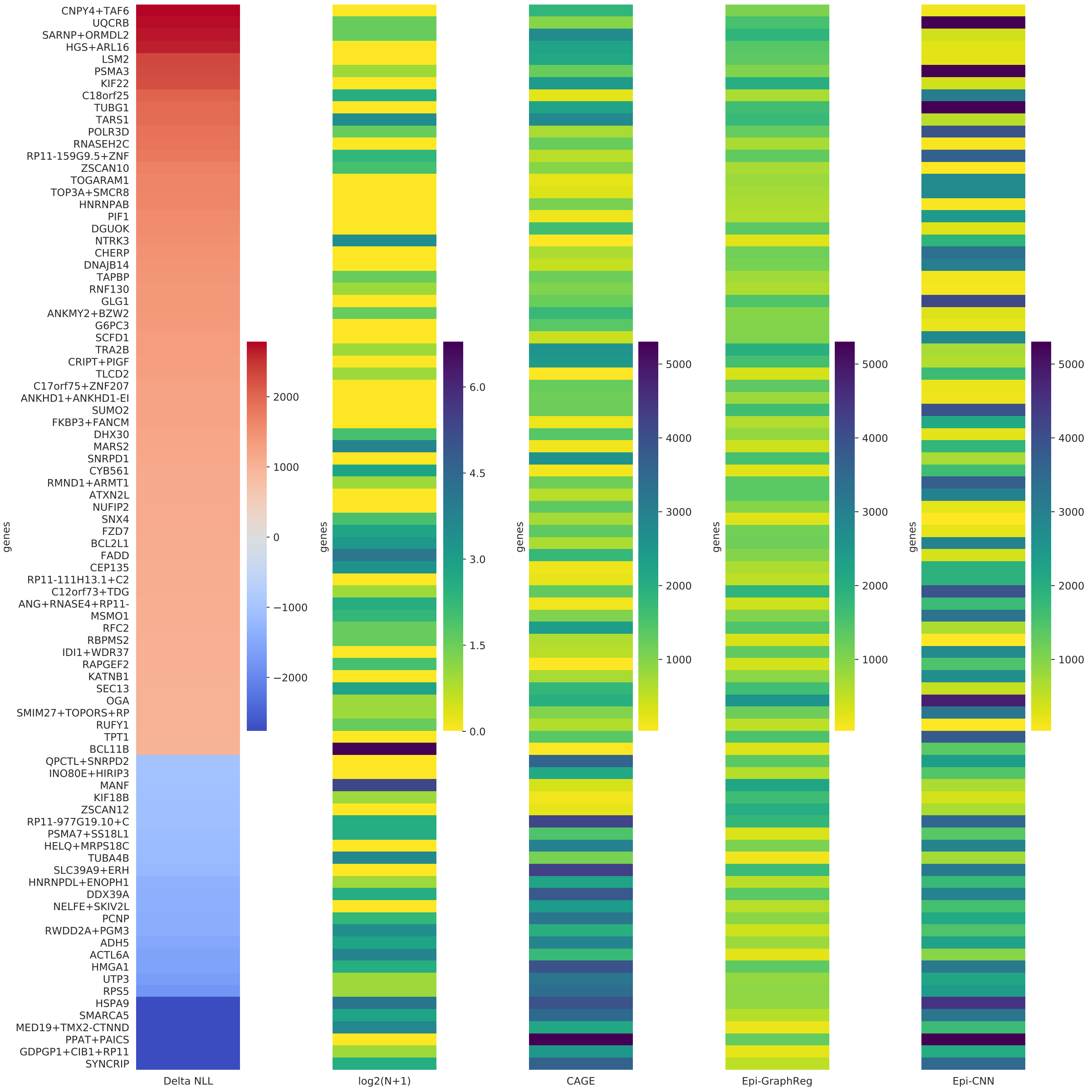


Figure S16: **hESC genes predicted accurately by either Epi-GraphReg or Epi-CNN and with significant difference in performance (Delta NLL)**. The results are for Micro-C (FDR=0.1).

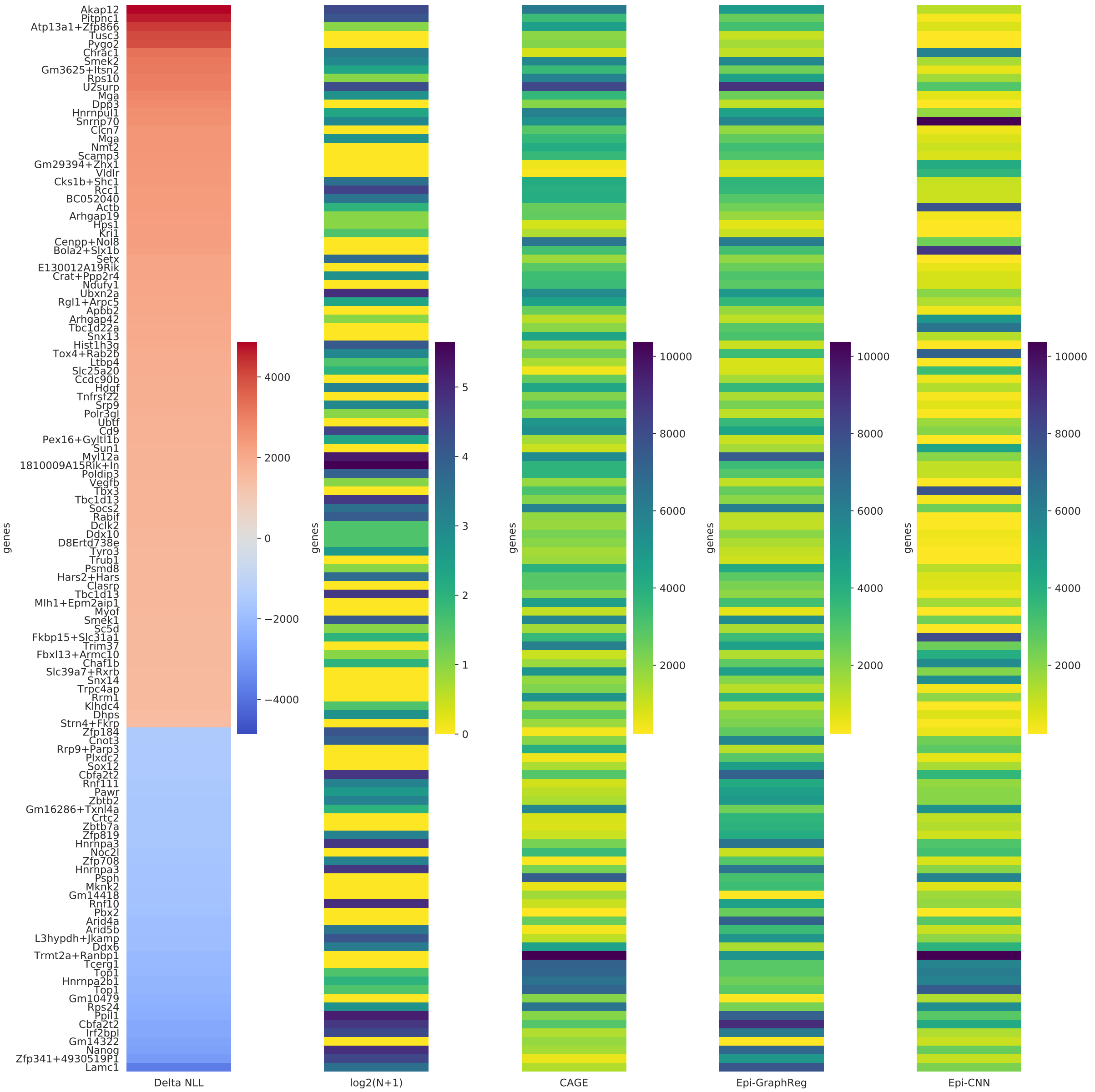


Figure S17: **mESC genes predicted accurately by either Epi-GraphReg or Epi-CNN and with significant difference in performance (Delta NLL)**. The results are for HiChIP (FDR=0.1).

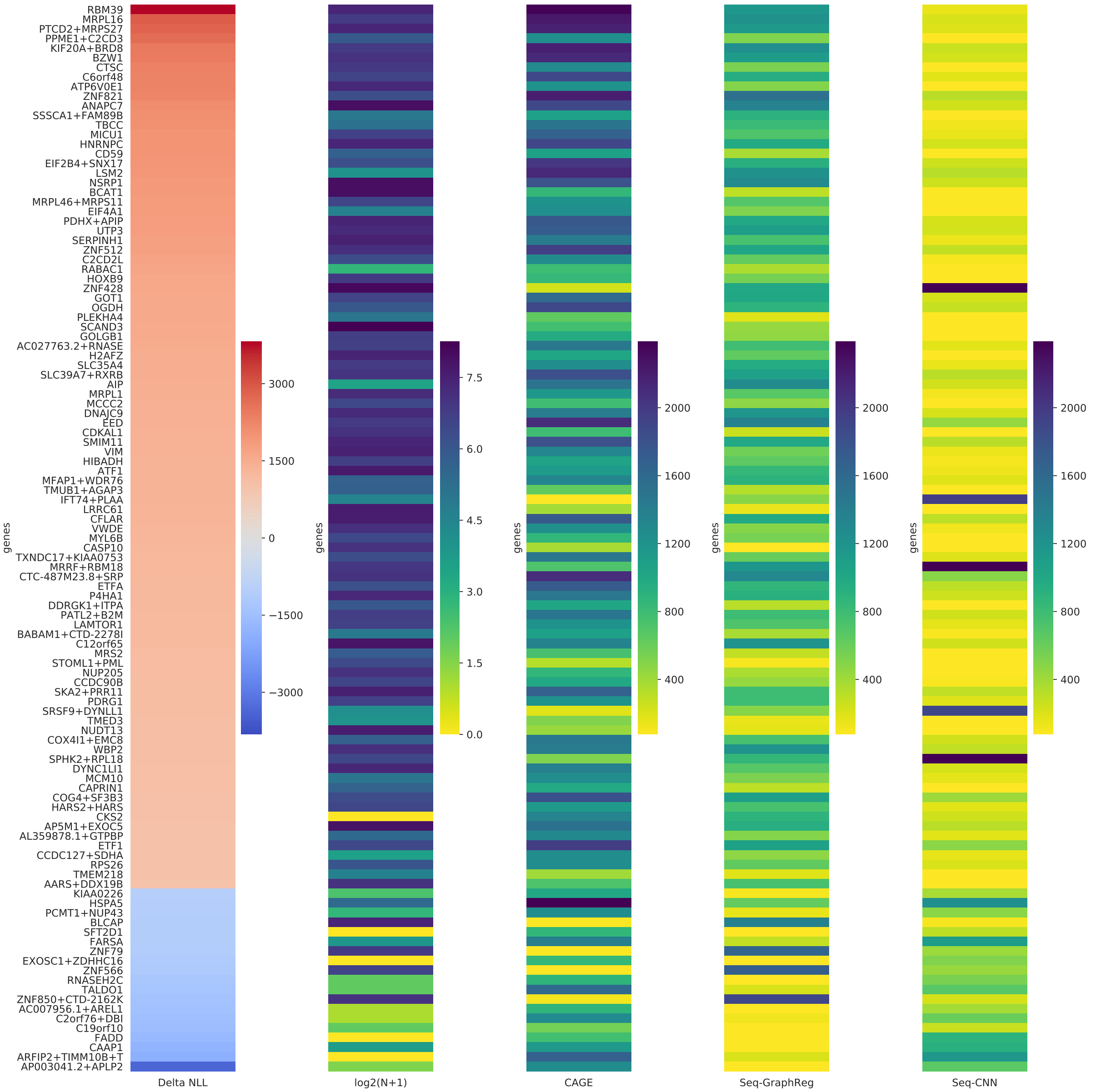


Figure S18: **K562 genes predicted accurately by either Seq-GraphReg or Seq-CNN and with significant difference in performance (Delta NLL)**. The results are for HiChIP (FDR=0.1) and separate training with no dilated layers.

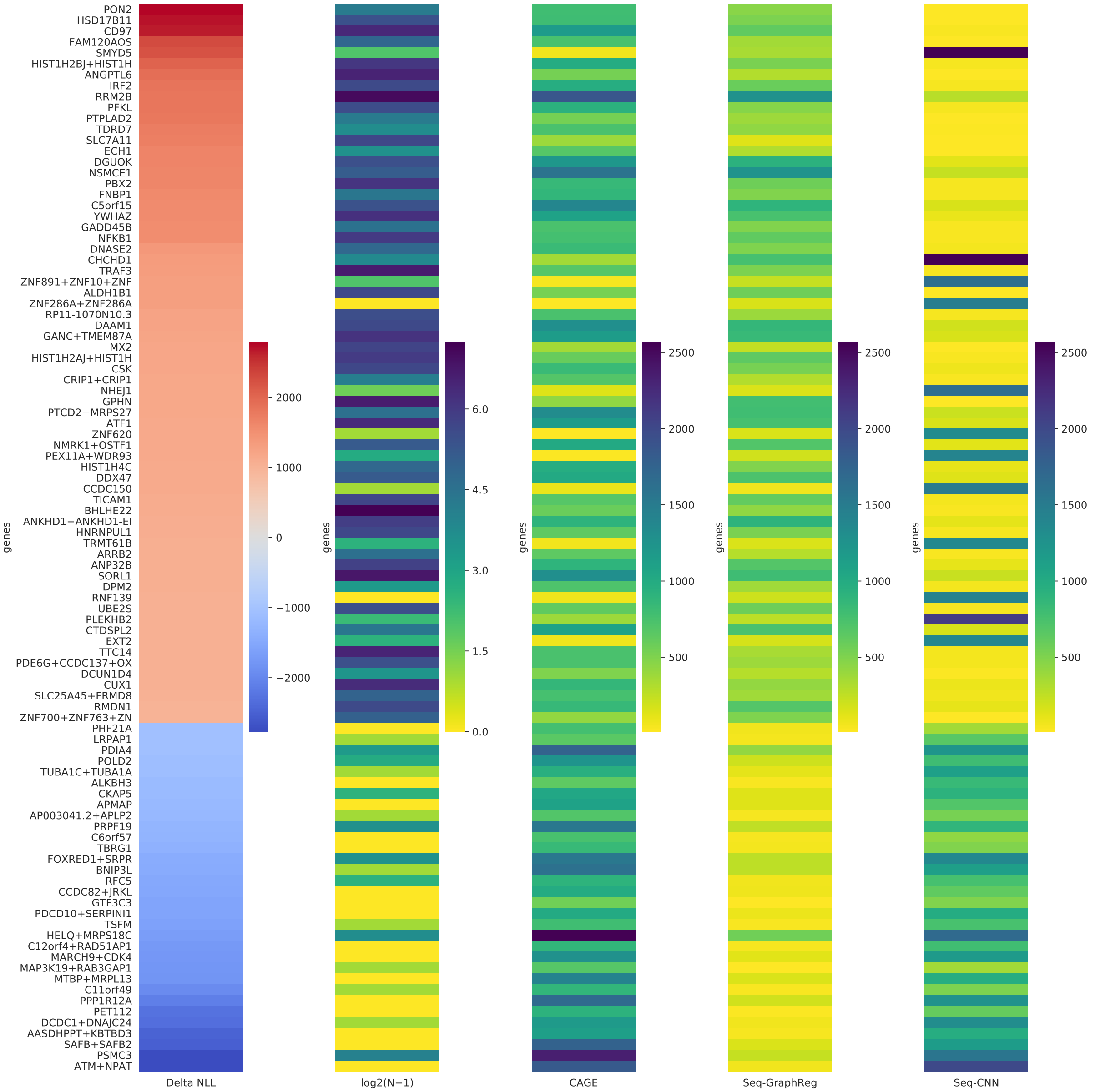


Figure S19: **GM12878 genes predicted accurately by either Seq-GraphReg or Seq-CNN and with significant difference in performance (Delta NLL)**. The results are for HiChIP (FDR=0.1) and end-to-end training.

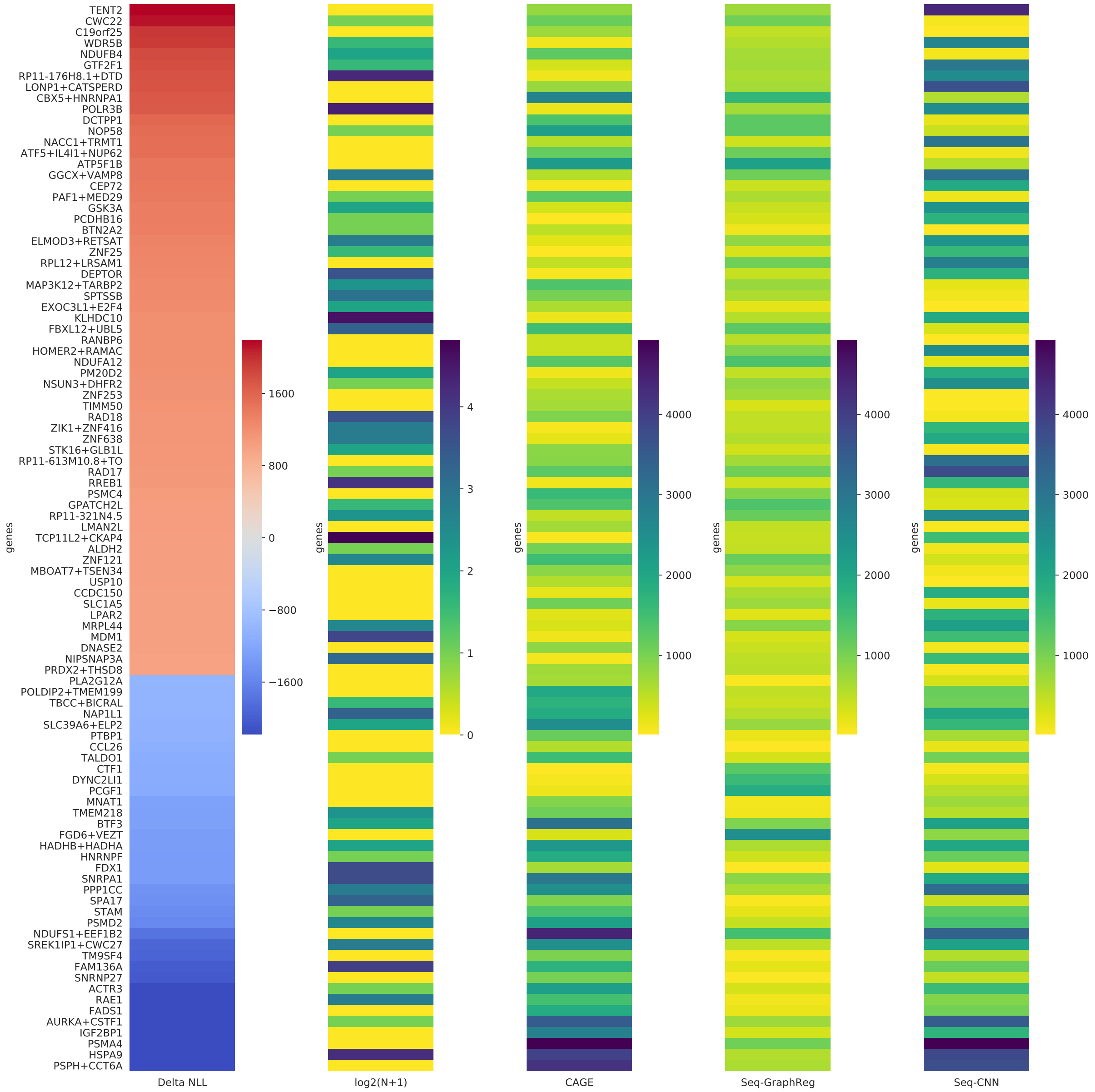


Figure S20: **hESC genes predicted accurately by either Seq-GraphReg or Seq-CNN and with significant difference in performance (Delta NLL)**. The results are for Micro-C (FDR=0.1) and end-to-end training.

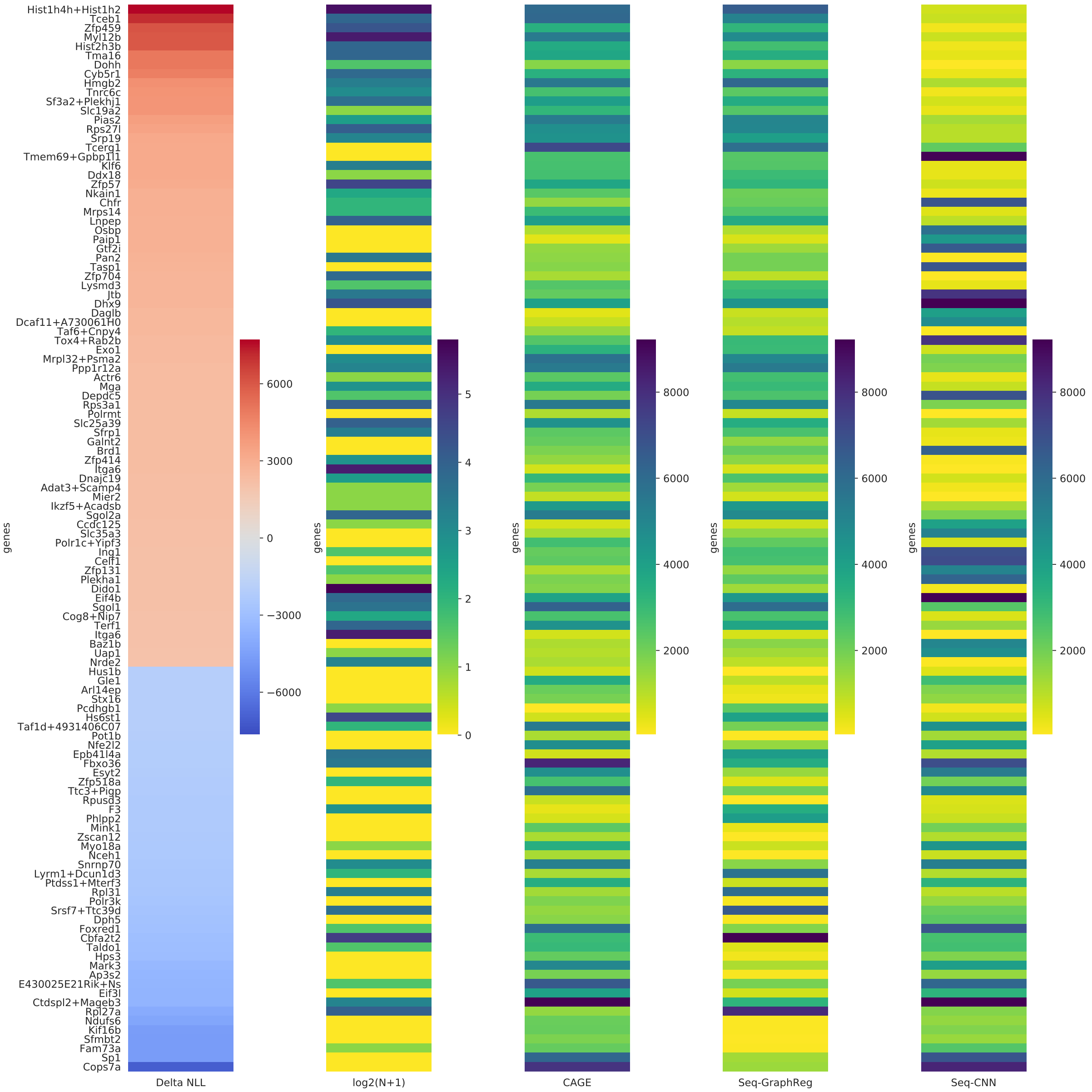


Figure S21: **mESC genes predicted accurately by either Seq-GraphReg or Seq-CNN and with significant difference in performance (Delta NLL)**. The results are for HiChIP (FDR=0.1) and end-to-end training.

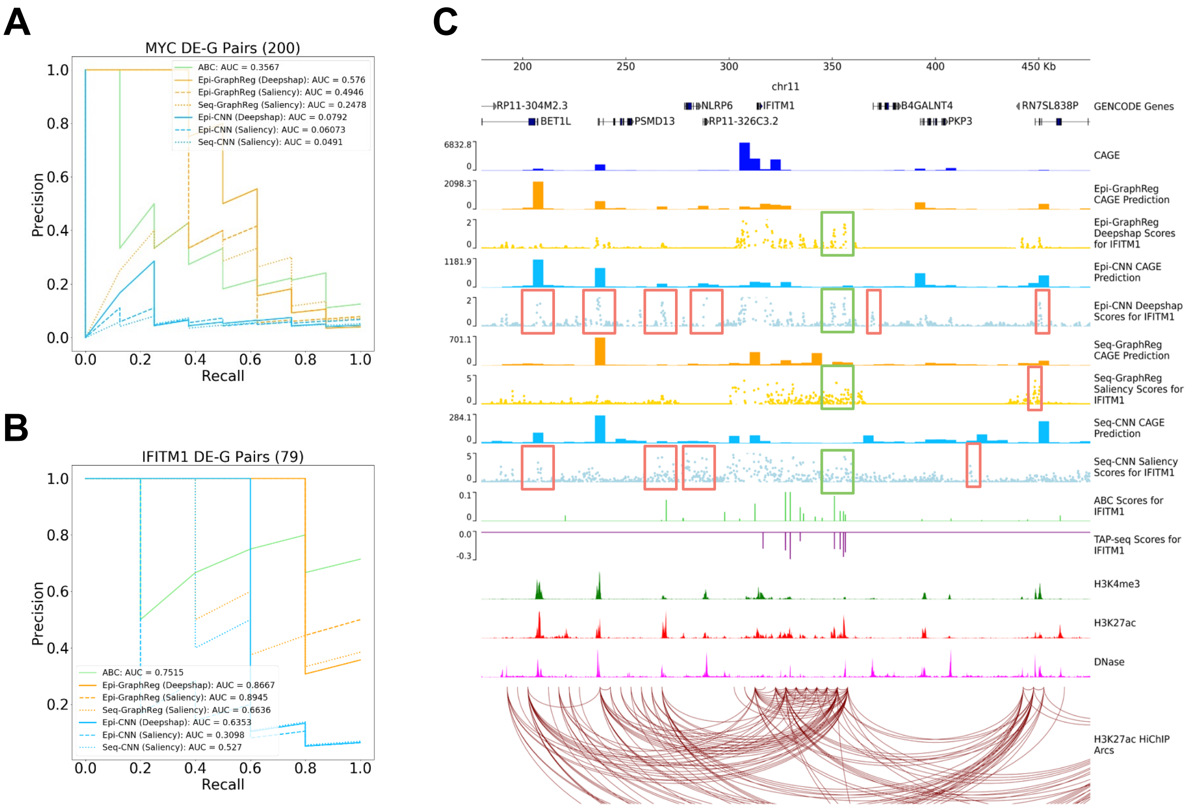


Figure S22: **GraphReg models more accurately identify functional enhancers of genes. A.** Precision-recall curves of the GraphReg, CNN, and ABC models for identifying enhancers of *MYC*. **B.** Precision-recall curves for the GraphReg, CNN, and ABC models for identifying functional enhancers of IFITM1. **C.** *IFITM1* locus (250Kb) in chr11 with epigenomic data, true CAGE, predicted CAGE using GraphReg and CNN models, HiChIP interaction graph, and the saliency maps of the GraphReg and CNN models, all in K562 cells. Experimental TAP-seq results and ABC values are also shown for *IFIM1*. Green and red boxes show the true positives and false positives, respectively. CNN models assign most of the nearby peaks as the true enhancers, leading to many false positives in promoter-proximal regions. GraphReg avoids this problem by explicitly attending to the likely regulatory regions based on HiChIP arcs in the graph.

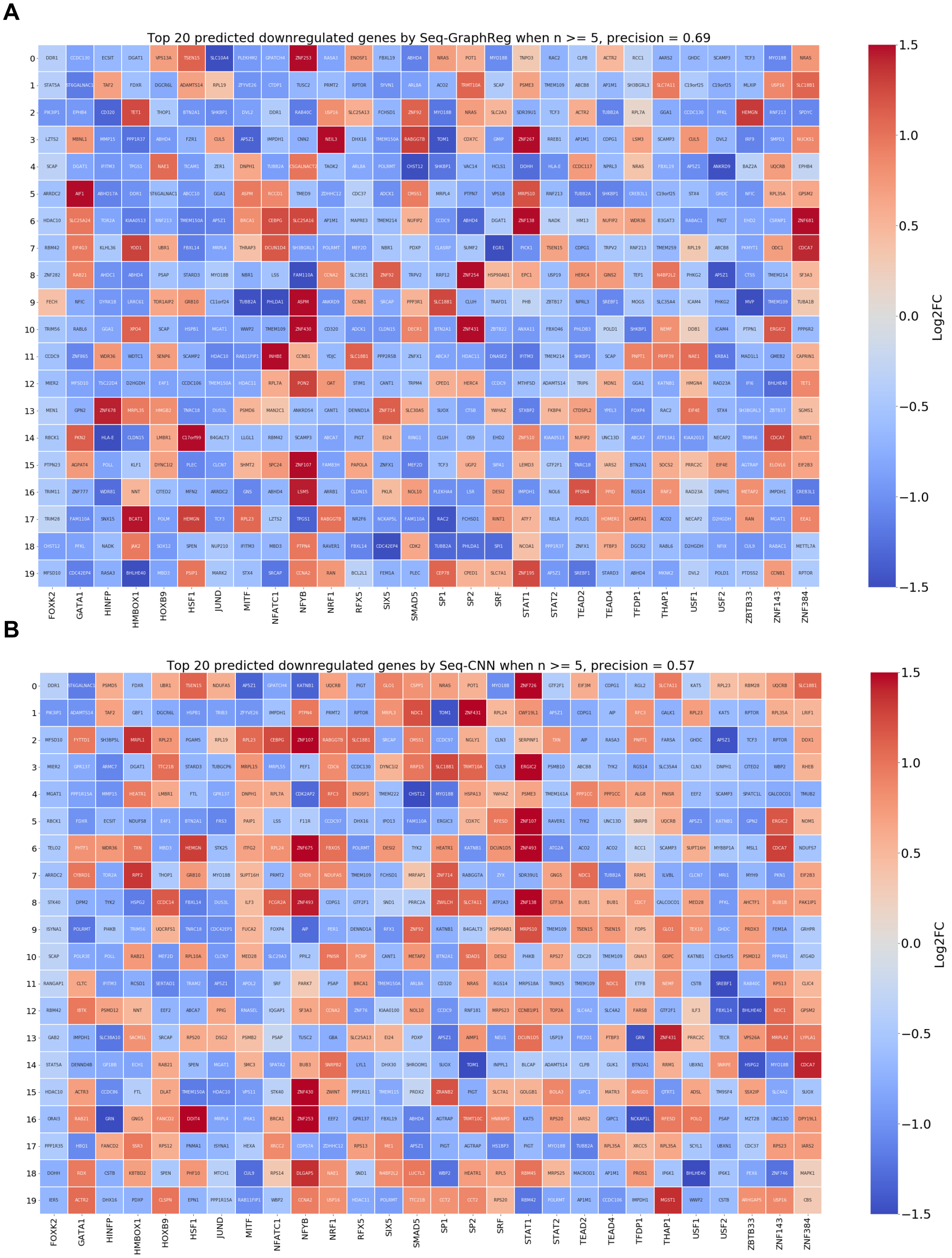


Figure S23: **Seq-GraphReg accurately predicts the regulatory effects of transcription factor knockouts by *in silico* motif ablation. A,B.** Top 20 predicted downregulated genes for each TF by the Seq-GraphReg and Seq-CNN models for $n \geq 5$, respectively. The genes are color-coded by the true logFC. All assessed genes are significantly differential ($p_{adj} < 0.05$). Predicted target genes with negative true logFC (blue colors) are correct predictions, those with positive true logFC (red colors) are false predictions. The precision of GraphReg model is 0.69 compared to 0.57 for the Seq-CNN model.
